# Gene regulation, phenotypic memory, and selection in fluctuating environments

**DOI:** 10.1101/2025.06.23.661119

**Authors:** Dan Pollack, Takashi Nozoe, Edo Kussell

## Abstract

Gene regulatory networks coordinate how microorganisms respond to environmental changes, and their proper functioning is critical for microbial survival and adaptation [1, 2]. While evolution of gene regulation has been extensively studied by comparative functional genomics [3, 4], the role of fluctuating environments in selecting for, shaping, and maintaining gene regulatory networks has not been elucidated. Laboratory evolution experiments find that under metabolic fluctuations, regulation of metabolic genes is often lost, leading to their constitutive expression [5–8]. To identify general properties of fluctuating environments that tune expression levels and select for gene regulation, we quantify the impact of regulation on fitness in strains with perturbed gene expression dynamics under metabolic fluctuations. We reveal that expression levels can be shaped by selection to enhance phenotypic memory of past environmental states, reducing the impact of gene induction lags on long-term population growth. By quantifying growth dynamics in a wide range of metabolic fluctuations, we identify catabolite symport flux as a key determinant of selection for gene regulation. By independently perturbing the ability to sense the environment and the control of expression levels, we discover sign epistasis between sensing and control in a gene regulatory network, and identify its molecular underpinnings. Due to this epistatic interaction, maintenance of sensing enhances the ability of evolution to tune gene expression levels in a fluctuating environment. Our work establishes a new basis for understanding how gene regulatory networks evolve in fluctuating environments.

## Introduction

Gene regulation functions to activate or repress gene expression when environments change, and to control expression levels when conditions remain constant. While these two distinct functions – sensing and control – are understood in molecular detail in different microbial model systems [2, 9–13], how fluctuating environments select for and maintain gene regulation during evolution has not been elucidated [1, 3]. The ability to sense and respond to fluctuations can be lost during episodes in which conditions remain constant for sufficiently long times, an outcome that has been observed in evolution experiments [5, 6, 8]. For example, *Escherichia coli* evolving in constant lactose conditions lose the ability to regulate the *lac* operon and express the *lac* genes constitutively, both *in vitro* [5] and *in vivo* [7]. Surprisingly, the same occurs when bacteria evolve in a periodic environment that alternates daily between lactose and glucose: mutations in the *lac* repressor (LacI) sweep to fixation and the evolved strains lose the ability to respond to lactose fluctuations [5, 14]. Indeed, activating and repressing metabolic genes repeatedly in a fluctuating environment could lead to recurring lag phases during gene induction (i.e. classic Monod lags) [15] which might select against gene regulation. However, the wild-type *lac* operon regulatory architecture is highly conserved in nature [16], indicating that natural selection acts to maintain gene regulation in this metabolic system due to mechanisms or constraints that are not currently understood [17].

In a fluctuating environment, bacterial fitness is determined by both exponential and non-exponential phases of growth, including lag and recovery phases [18]. To optimize growth, cells ‘remember’ their past metabolic states to minimize or avoid repeated gene induction lags, a behavior known as phenotypic memory [19, 20]. Phenotypic memory in the *lac* operon relies on a simple molecular mechanism: the inheritance from mother to daughter cells of stable metabolic proteins, the enzyme β-galactosidase (LacZ) and the lactose permease (LacY), which together enable growth on lactose. Phenotypic memory was shown to eliminate gene induction lags up to 4 generations after lactose was first encountered, or to shorten lag durations up to 10 generations after initial exposure [19]. Modulating the expression levels of the *lac* operon, which are controlled by both the repressor (LacI) and the cyclic-AMP receptor protein (Crp), is thus predicted to alter the duration of memory. While previous studies have considered how control of gene expression levels impacts bacterial fitness in constant environments [21–25], the molecular mechanism of phenotypic memory suggests that gene expression levels may in fact be tuned to provide memory to future generations in a fluctuating environment. Yet whether and how strongly selection may act to enhance phenotypic memory, and thereby to reduce the deleterious effect of gene induction lags on long-term growth, has not been explored.

Taking the above to its logical extreme, memory of arbitrary duration – corresponding to constitutive expression or the absence of gene regulation – is expected to eliminate gene induction lags altogether. Identifying fluctuating conditions that select against constitutive expression is therefore critical for understanding how gene regulation is maintained in nature. Early work on bacterial physiology established that constitutive *lac* expression can result in growth arrest or even cell death upon initial exposure to saturating concentrations of lactose when cells are grown on different carbon sources [26, 27]. Such behavior in constitutive *lac* mutants, which we will refer to as lactose-induced growth arrest, was later shown to result from wasteful LacY activity in the presence of excess lactose, which diffuses the proton motive force (PMF) during lactose import and reduces ATP production [28, 29]. Depending on the specific media and genetic background, constitutive *lac* cells may recover from such arrest at different rates [26, 28–30] or be killed [27, 28]. In contrast, wild-type (i.e. regulated) *lac* expression is not susceptible to lactose-induced growth arrest, suggesting that specific metabolic transitions could select for regulation. Alternatively, while attenuated by phenotypic memory, wild-type regulation is still susceptible to gene induction lags. Fluctuating environments that enhance gene induction lags while minimizing lactose-induced growth arrest could thus select for constitutive expression.

Given the intricate relationships between gene expression dynamics, phenotypic memory, and bacterial fitness in a fluctuating environment, understanding the adaptive advantage of gene regulation requires analyzing the molecular underpinnings of growth rates and lag phases, as well as their dependence on environmental conditions and fluctuation timescales. Importantly, these must be assessed in well-defined mutational contexts that perturb distinct features of a gene regulatory network, e.g. abolishing the ability to sense and respond, altering maximal expression levels, reducing the speed of induction, etc. Here, we systematically analyze these relationships using targeted perturbations of gene regulation in the *lac* operon. Using bulk population and microfluidic growth rate measurements, we determine which fluctuating environments select for or against gene regulation. By competing strains with perturbed gene regulation and altered levels of phenotypic memory, and tracking strain frequency dynamics using barcode sequencing in a fluctuating environment, we measure the strength of selection acting on gene regulation and tuning expression levels. Our findings establish fundamental principles of how evolution shapes gene regulation and memory in bacteria, and have broader implications in studies of microbial communities [31], synthetic biology [2], metabolism [12], and metabolic engineering [32].

## Results

### Selection for phenotypic memory

Wild-type *E. coli* growing under periodic glucose-lactose fluctuations in a microfluidic device exhibit a gene induction lag upon initial exposure to lactose **(Fig. 1A; Movie S1)**. Due to phenotypic memory resulting from inheritance of stable Lac proteins, subsequent exposures to lactose do not result in lag phases **(Fig. 1A)**, as residual Lac protein levels are sufficient to support growth [19]. To determine conditions that select for phenotypic memory, we used a library of *E. coli* strains in which we destabilized the β-galactosidase enzyme with ssrA degradation tags spanning a range of degradation rates [33]. Despite reduction of β-galactosidase expression levels by up to 30% during steady-state growth in these *lacZssrA* strains **(Table S1)**, no significant growth rate differences were observed in minimal lactose media or other conditions **(Fig. 1B** and **Table S1)**. We quantified lag phase duration of each strain in glucose-to-lactose transitions, and found that strains with the lowest β-galactosidase expression levels (*NYNY* and *GSNY*) had the longest lag phase durations **(Fig. 1C)**. We therefore predicted that selection would not act on gene expression levels in a constant lactose environment, but could act against lower expression in fluctuating glucose-lactose conditions due to the presence of increased lag phases. We propagated the mixture of *lacZssrA* strains as a single batch culture in minimal media supplemented with either glucose or lactose as the only carbon source. Cultures were grown in either constant conditions or daily fluctuations between the two carbon sources, and samples were assayed at regular intervals to determine relative strain frequencies **(Fig. 1D)**. We initially ran a short pilot experiment lasting 5 days, and found that cultures grown in daily fluctuations between glucose and lactose exhibited selection against strains with the lowest expression levels, while cultures grown in constant conditions showed no significant fitness differences among the strains **(Fig. S1)**. We then repeated the experiment, expanding its duration to 14 days, and running separate experiments at three different daily dilution factors **(Fig. 1E** and **Figs. S2-S4)**. In each case, the experiments revealed that daily fluctuations between glucose and lactose select for higher *lac* expression levels. Measurement of selection coefficients revealed small (∼1%), sustained and statistically significant fitness differences among the strains in the fluctuating environment **(Fig. 1F)**. Modeling of expression levels and induction kinetics recapitulated the overall small magnitude of selection coefficients **(Fig. S4C,F)**, and predicted that the strength of selection for memory decreases with number of generations spent in each environment **(Fig. 1G)**. Consistent with the model, the highest selection coefficients were measured for the *No Deg* strain in the 1:16 daily cell dilution corresponding to the highest level of memory and the lowest number of generations in each condition **(Fig. 1G)**. Due to the larger number of generations in experiments with stronger dilution factors, beneficial mutations unrelated to *lac* expression were found to increase in frequency in the second half of several experiments **(Figs. S2-S4** and **Supplementary Text)**. In such cases, selection coefficients for *lac* expression levels were inferred from the initial part of the experiment up to the point at which a selective sweep was observed to begin. Together, these results support our hypothesis that selection favors higher β-galactosidase expression levels in fluctuating environments, increasing the duration of phenotypic memory, and reducing growth lags that result from gene induction kinetics.

**Fig. 1.**
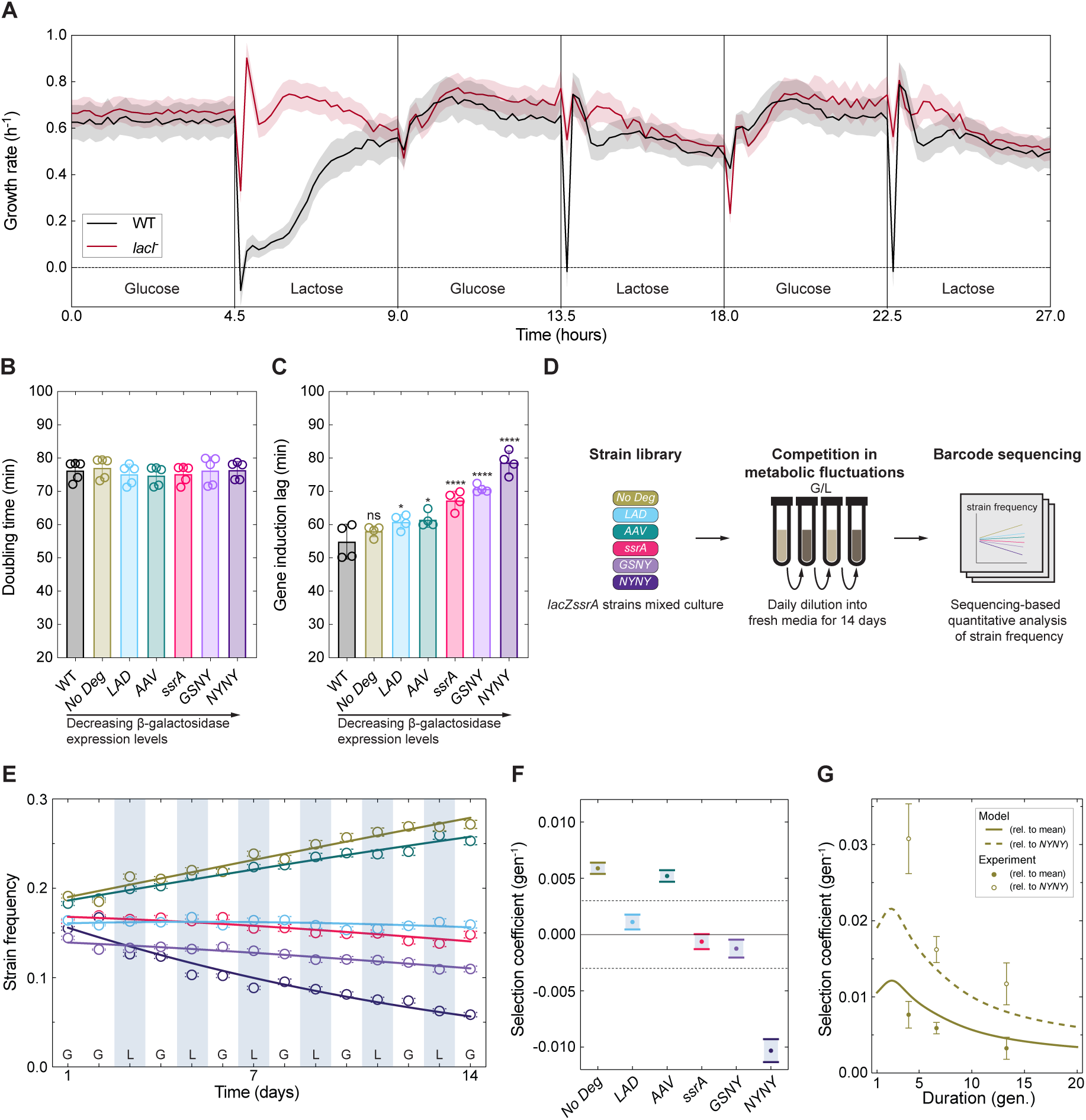
Selection for phenotypic memory in a fluctuating environment. **(A)** Growth rate measurements of WT and *lacI^−^* strains in periodic glucose-lactose fluctuations using microfluidics. Cells were grown in minimal media with 0.4% lactose or 0.2% glucose as carbon source, and supplemented with 1% casamino acids. Mean growth rate of cells in microfluidic chambers is shown (solid line) ± standard error of the mean (SEM) (shaded envelope). See also Movies S1,2. **(B)** Doubling time measured for WT *E. coli* and *lacZssrA* strains grown in lactose minimal media. Strains are ordered according to β-galactosidase expression levels during induction in lactose. Bars and error bars indicate mean ± std. dev. (SD) over biological replicates (open circles, n = 5); one-way ANOVA, *p*-value = 0.89. See also Table S1 for additional growth rate measurements and β-galactosidase activity levels. **(C)** Duration of lag phase upon glucose-to-lactose transition. One-way ANOVA with Dunnett multiple comparisons test of each strain to the WT was used for statistical analysis. Statistically significant differences are indicated (\*\*\*\**p<*1e-4, \**p<*0.05, ns = *p*>0.05). **(D)** Schematic of mixed-culture competition assay. The library of *lacZssrA* strains in equal ratios was inoculated at *t* = 1 and propagated in 9 parallel experiments for 14 days. Strains were sampled and diluted (1:16, 1:100, or 1:10,000) into fresh media alternating glucose/lactose (G/L) every 24 h. Controls were grown for the same duration and dilutions in either constant glucose or lactose media. DNA was extracted from each sample and the 3’ end of the *lacZ* gene was amplified by PCR. Pooled samples were sequenced and read counts were used to determine strain frequencies. **(E)** Frequency of *lacZssrA* strains over two weeks grown in alternating G/L with 1:100 daily dilution. Relative exponential growth model was fit (solid lines) to infer selection coefficients. Error bars indicate ± SEM based on a multinomial distribution of read counts. **(F)** Relative selection coefficients (points) and 95% confidence intervals (bars) are shown for each strain inferred from model fitting in E. Dashed lines indicate upper and lower bound of coefficient confidence intervals inferred from competition in constant glucose. See also Fig. S1-S4 and supplementary text for further details. **(G)** Predicted dependence of relative selection coefficients on number of generations between environmental changes. Model predictions (curves) and measurements (points) of selection coefficients are shown for the *No Deg* strain relative to either the *NYNY* strain or the mean over all 6 strains. See also Fig. S4C,F for predicted competitive dynamics of the *lacZssrA* strain library and relative selection coefficients.

### Impact of gene regulation on fitness in constant and fluctuating environments

We deleted the *lacI* gene (*lacI^−^*) in the *E. coli* chromosome to yield constitutive *lac* expression and compared *lac* expression levels, doubling times, and bioenergetics of WT and *lacI^−^* strains in various growth conditions **(Fig. 2A,B; Fig. S5** and **Table S2)**. We measured the instantaneous growth rate of the *lacI^−^* strain in periodic glucose-lactose fluctuations using microfluidics **(Fig. 1A; Movie S2)**. Unlike WT, the *lacI^−^* strain grew continuously through the first transition to lactose without exhibiting a gene induction lag. We quantified the fitness cost of constitutive expression in a constant environment by comparing doubling times of WT and *lacI^−^*in a panel of carbon sources. Despite the substantial increase in *lac* expression levels resulting from increasing levels of cAMP as intracellular glucose levels decline in poorer media [11] **(Fig. 2A)**, we did not detect a difference in doubling times between WT and *lacI^−^* in any carbon source that we tested **(Fig. 2B; Fig. S5)**. For example, in succinate the *lacI^−^* strain over-expresses the *lac* operon by 4.5 times relative to normal induction levels in lactose **(Fig. 2A; Table S2)**. The WT strain produces no detectible level of *lac* expression in any condition except lactose **(Fig. 2A)**, yet its doubling time is indistinguishable from that of *lacI^−^*in each of the 8 carbon sources tested **(Fig. 2B)**. We tested if activity of the *lac* permease impacted growth by exposing cells to the non-metabolizable inducer isopropyl β-D-1-thiogalactopyranoside (IPTG) in each condition. Consistent with previous findings, we found longer doubling times in the presence of IPTG [26, 27, 29, 30], which increased with higher *lac* expression levels [25] **(Fig. S5A,B)**. However, we found that WT and *lacI^−^* strains grew identically in all conditions **(Fig. S5A,B)**. We found statistically significant differences in the oxygen consumption rate (OCR) with and without IPTG, confirming that aerobic respiration was impacted by LacY activity, but we did not find differences in OCR between WT and *lacI^−^* strains **(Fig. S5C)**. Our results demonstrate that despite substantial differences in doubling times and expression levels across conditions, in a constant environment constitutive *lac* expression does not incur a fitness cost relative to wild-type expression.

**Fig. 2.**
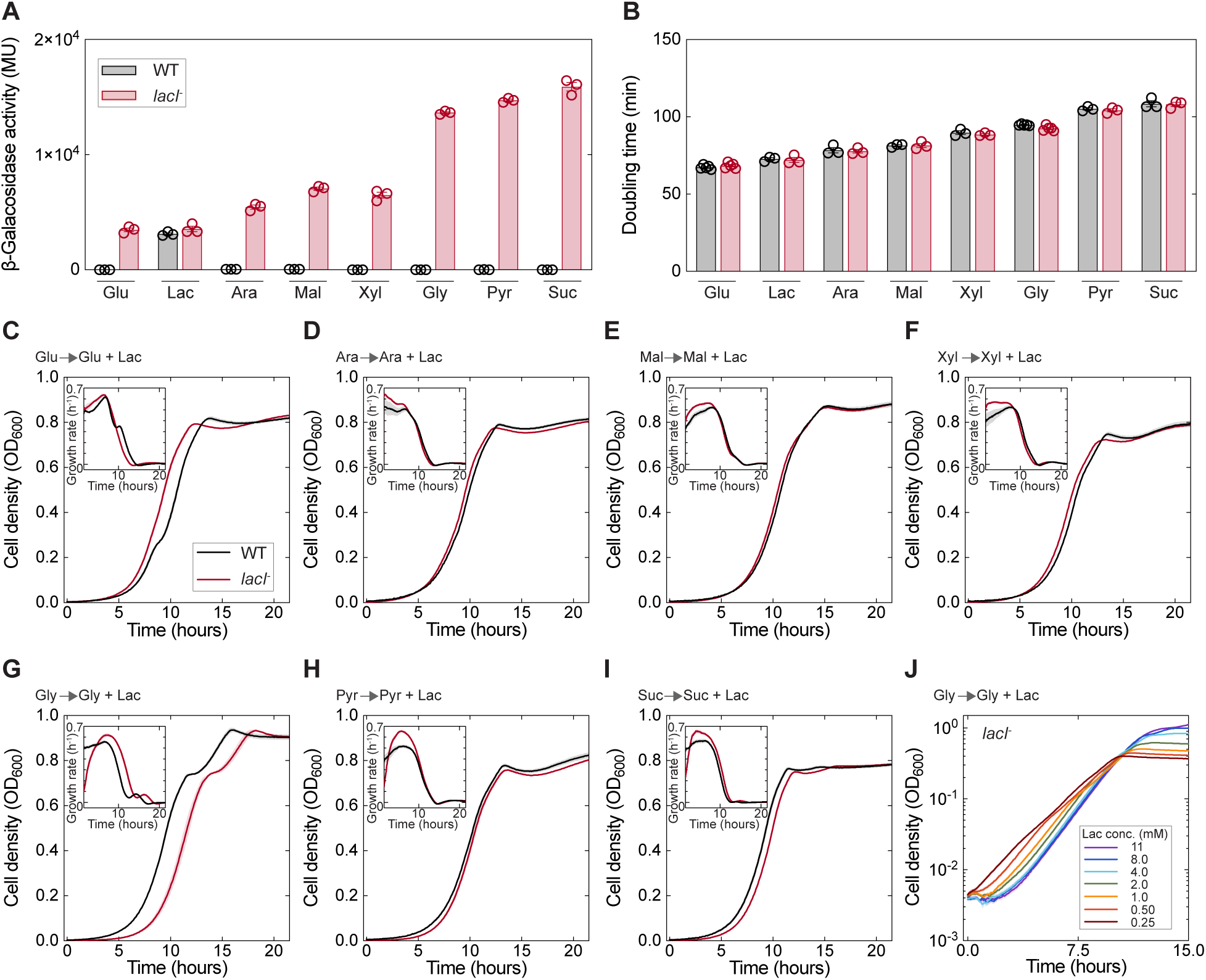
Growth dynamics and expression levels with and without gene regulation. **(A)** β-Galactosidase activity levels in Miller units (MU) are shown for WT and *lacI^−^* strains grown in minimal media supplemented with glucose, lactose, arabinose, maltose, xylose, glycerol, pyruvate, or succinate. Bars indicate mean ± SD (n = 3). See also Fig. S5A for additional β-galactosidase activity in each carbon source with IPTG and statistical analysis. **(B)** Doubling time measured for WT and *lacI^−^*strains in the same conditions as panel A. Individual biological replicates (n ≥ 3) are shown in circles. See also Fig. S5B for additional growth measurements in each carbon source with IPTG and statistical analysis. **(C-I)** Diauxic growth dynamics of WT and *lacI^−^* strains grown separately in a mixture (1:2 v/v) of the specified carbon source and lactose minimal media. OD_600_ and instantaneous growth rates (insets) shown as mean (solid line) ± SEM (shaded envelope) over biological replicates (n = 3). Cells were grown to exponential phase in the specified carbon source and diluted to an OD_600_ = 0.01 at *t* = 0. See also Fig. S6A-G for diauxic growth dynamics in a mixture (2:1 v/v). **(J)** *lacI^−^*strain grown in mixture (1:2 v/v) of glycerol and lactose minimal media. Cells were grown to exponential phase in glycerol and diluted to an OD_600_ = 0.01 at *t* = 0. Legend shows the molarity of lactose for each growth condition.

We measured the instantaneous growth rate of WT and *lacI^−^*during transitions from a single carbon source into diauxic growth on a mixture of two carbon sources [15]. In such conditions, wild-type *E. coli* consume carbon sources sequentially, due to regulation by the carbon catabolite repression (CCR) mechanisms, with glucose as the preferred carbon source [11, 13]. We grew WT and *lacI^−^* strains in one of seven different carbon sources and then transferred cells to a mixture of the same carbon source and lactose **(Fig. 2C-I; Fig. S6A-G)**. When transferred from glucose to diauxic growth on glucose and lactose **(Fig. 2C; Fig. S6A)**, both strains were able to grow immediately upon transfer (at *t* = 0), which is expected given the availability of glucose. The *lacI^−^* strain exhibited a small growth advantage in the first 5 hours of the experiment **(Fig 2C**, inset**; Fig. S6A,** inset). Similarly, *lacI^−^* exhibited faster initial growth than WT in lactose diauxic conditions with arabinose, xylose, or maltose **(Fig. 2C-F; Fig. S6B-D)**. In contrast, when transferred from glycerol to diauxic growth on glycerol and lactose **(Fig. 2G)**, the *lacI^−^* strain exhibited a prolonged lag phase while WT grew rapidly from the outset at *t* = 0 **(Fig. 2G**, inset**; Fig. S6E,** inset). Similarly, a prolonged initial lag was observed for *lacI^−^*but not WT upon transfer into lactose diauxie with pyruvate or succinate **(Fig. 2H,I**, insets**; Fig. S6F,G,** insets).

To determine whether excessive transport of lactose was directly responsible for the lags observed in the *lacI^−^* strain **(Fig. 2G-I)**, we grew *lacI^−^*in varying concentrations of lactose to modulate the flux of lactose into cells after initial growth in glycerol. We found that lag duration decreased with lower lactose concentrations, reaching a point where lags were eliminated, and exponential growth took place from the outset **(Fig. 2J)**. Thus, the lags observed in the *lacI^−^* strain exhibit similar characteristics to lactose-induced growth arrest [26]. Further, we did not observe killing when plating on lactose, as previously found [27, 28]; cells remained viable during the lag phase and cell counts increased during the recovery phase, similar to previous observations [26] **(Fig. S6H,I)**. Given that a lag is only observed above a threshold concentration of lactose, and becomes more pronounced with increasing concentration, hereafter we refer to this type of lag phase as a ‘catabolite hyperflux lag’. Since lactose flux is limited by the number of permease molecules, the duration of the lag is expected to approach a maximum once the permease is working at saturation, i.e. for sufficiently high concentrations of lactose, consistent with the data **(Fig. 2J)**. Working at a saturating concentration of lactose and modulating *lac* permease levels by growing the *lacI^−^* strain in different carbon sources, we estimate the threshold *lac* expression level required for catabolite hyperflux lag is ∼3 times normal expression levels in lactose **(Fig. 2A** and **Table S2)**.

### Selection for gene regulation

Given the advantage of *lacI^−^* during metabolic transitions below the catabolite hyperflux threshold **(Fig. 2C-F)**, and the advantage of WT above it **(Fig. 2G-I)**, we hypothesized that *lac* regulation could be selected either for or against in fluctuating lactose diauxic growth conditions. We tested this hypothesis by competition experiments, in which we co-cultured WT, *lacI^−^*, and *NYNY* strains and subjected these strains to daily fluctuations between growth in a single carbon source (glucose or glycerol) and diauxic growth conditions with lactose (see Methods). In periodic glucose and lactose diauxie, we observed a continuous increase in *lacI^−^*frequency (selection coefficient ∼0.7%) while WT remained unchanged and *NYNY* decreased from its starting frequency **(Fig. 3A,B)**. This confirms that constitutive expression is beneficial in conditions for which *lac* expression levels remain below the catabolite hyperflux threshold **(Fig. 2C)**. Additionally, the gradual decrease of *NYNY* recapitulates our previous observation regarding the benefit of phenotypic memory in glucose-lactose fluctuations (compare **Fig. 1E)**. In periodic glycerol and lactose diauxie, we observed extinction of the *lacI^−^* strain by day 11 **(Fig. 3C)**, while the WT strain frequency increased relative to *lacI^−^* with a selection coefficient of ∼5% **(Fig. 3D)**. This demonstrates the advantage of gene regulation in conditions where expression levels increase above the catabolite hyperflux threshold **(Fig. 2G)**. Moreover, once the frequency of *lacI^−^*had decayed, *NYNY* began to gradually decrease relative to WT **(Fig. 3C)**. In both control conditions, all three strains were maintained at similar frequencies throughout the experiment **(Fig. S7)**. These results establish that fluctuating lactose diauxie using a poor carbon source selects for gene regulation in the *lac* operon, while simultaneously selecting against lower *lac* expression levels that decrease phenotypic memory.

**Fig. 3.**
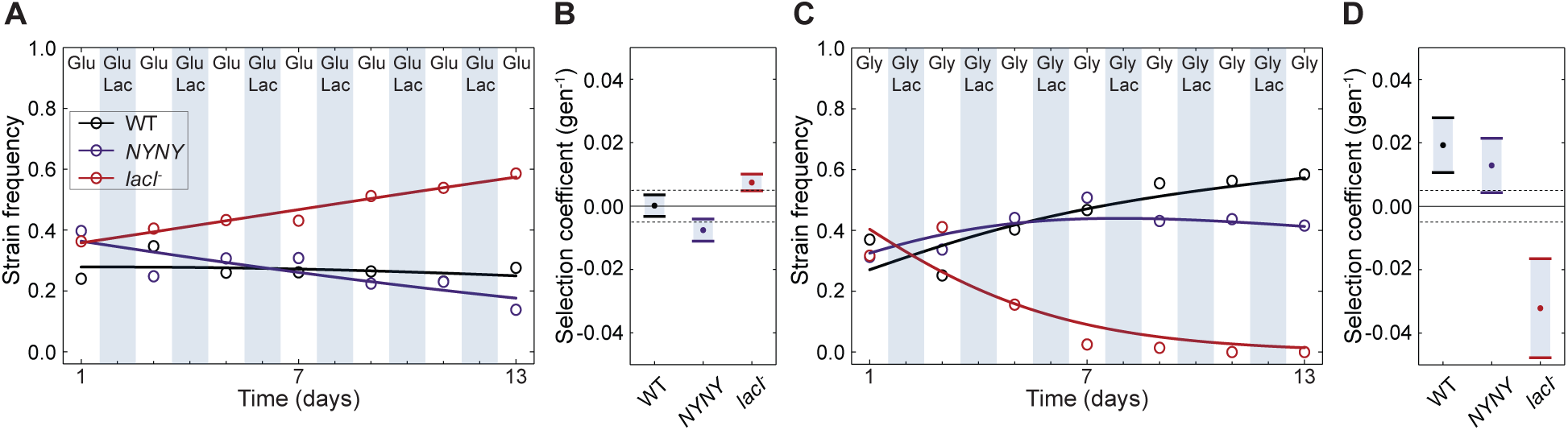
Selection for or against gene regulation in fluctuating diauxic conditions. Competition experiments in diauxic fluctuating environments. A mixed culture of WT, *lacI^−^*, and *NYNY* strains combined in equal ratios at *t* = 1 based on normalized OD_600_ was grown in a periodic environment alternating between glucose-only media and glucose and lactose diauxie, shown in panel **(A)**; or between glycerol-only media and glycerol and lactose diauxie, shown in panel **(C)**. Relative exponential growth model was fit (solid lines) to infer selection coefficients. **(B,D)** Relative selection coefficients (points) and 95% confidence intervals (bars) are shown for each strain inferred from model fitting in A,C respectively. Dashed lines indicate upper and lower bound of confidence intervals inferred from competition in constant glycerol. See also Fig. S7 for controls of competition experiments.

### Sign epistasis between sensing and control of expression in a gene regulatory network

To investigate how presence or absence of sensing interacts with genetic mutations that control expression levels, in both WT and *lacI^−^*strains we edited the UP element (-45 region) of the *lac* promoter, which impacts Crp-dependent activation [34–37], generating a library of strains with altered *lac* expression levels spanning 50% - 150% WT levels **(Table S3)**. For the unregulated strains (*lacI^−^_UE_*), a transition to glucose and lactose diauxie favored strains with higher expression levels, which reached higher OD values sooner **(Fig. 4A** and **Fig. S8A)**. In contrast, upon transition to glycerol and lactose diauxie, *lacI^−^_UE_* strains exhibited catabolite hyperflux lags, which favored strains with lower expression levels **(Fig. 4B** and **Fig. S8B)**. For the regulated strains (WT*_UE_*), both diauxic growth conditions favored strains with higher expression levels. In glucose and lactose diauxie, WT*_UE_* strains exhibited a diauxic lag phase at approximately *t* = 10 h when glucose is depleted and *lac* genes are induced, and strains with the highest expression levels had the shortest diauxic lags **(Fig. 4D** and **Fig. S7D)**. Similarly, in glycerol and lactose diauxie, where lactose is consumed first, WT*_UE_* strains with the highest expression levels exhibit a shorter gene induction lag and reached higher OD values sooner **(Fig. 4E,F** and **Fig. S8E,F)**. Comparing WT*_UE_* and *lacI^−^_UE_* strains, we find that variation in the lag and recovery time was significantly larger in the absence of regulation **(Fig. 4C,F; Fig. S8C,F)**. This demonstrates that the *lac* repressor acts to buffer the deleterious effects of genetic variation revealed in the *lacI^−^_UE_* library. Not only does the repressor reduce variation in lag durations which impact fitness, its presence changes the sign of fitness differences **(Fig. 4C,F)**. This genetic interaction, known as sign epistasis [17], between mutations impacting sensing and control of expression, may enhance the ability of gene regulatory networks to adapt as discussed below.

**Fig. 4.**
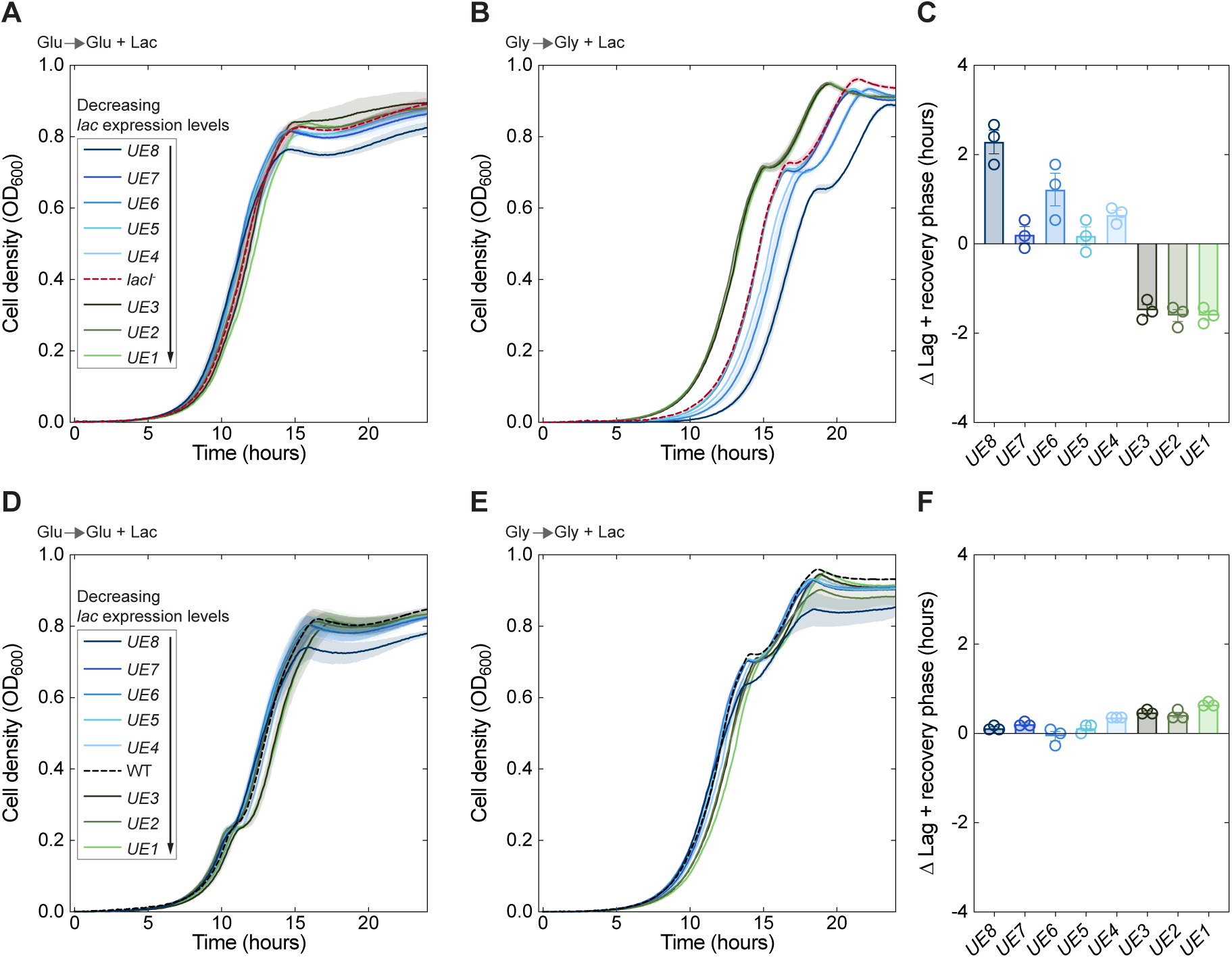
Epistatic interaction between sensing and control of gene expression. Diauxic growth dynamics of the UP element (UE) strain library in *lacI^−^***(A**,**B)** and WT **(D**,**E)** backgrounds. Each strain was grown separately in a mixture (1:2 v/v) of glucose (A,D) or glycerol (B,E) and lactose minimal media. Cells were grown overnight in glucose (A,D) or glycerol (B,E) and diluted to an OD_600_ = 0.01 at *t* = 0. Legend indicates strains ordered by *lac* operon expression levels. Curves show mean OD_600_ measurements taken every 5 minutes over biological replicates (n = 3); ± SEM is shown in shaded envelope. **(C,F)** Difference (Δ) in duration of lag and recovery phase during diauxic growth on glycerol and lactose (B,E) between each *lacI^−^_UE_* strain and *lacI^−^* (C) or between each WT*_UE_* strain and WT (F). Time to OD_600_ = 0.02 is shown measuring catabolite hyperflux lag and recovery (C) and gene induction lag and recovery (F). Variance of lag and recovery phase durations was calculated for each plot, yielding Var(Δ) = 2.36 h^2^ (C) or 0.05 h^2^ (F), which were found to be significantly different (F-test, *p*-value = 4.0e-14). Spearman’s correlation coefficient (ρ) was calculated between Δ values and *lac* expression levels over UE strains, ρ = -0.86, *p*-value = 0.006 (C) and ρ = 0.78, *p*-value = 0.02 (F). See also Fig. S8 for diauxic growth dynamics in a mixture (2:1 v/v) and Table S3 for UP-element strains’ modified sequence, β-galactosidase activity levels, and growth rates.

## Discussion

Our investigation provides a systematic basis for understanding the relation between bacterial physiology and the selective constraints that shape the two principal functions of a gene regulatory circuit – environmental sensing and control of expression levels. Mutations that disrupt gene regulation may impact both functions in different ways depending on environmental conditions. We performed experiments in constant and fluctuating environments to assess sensing and control in different genetic backgrounds that perturb regulation of the *lac* genes.

In constant conditions, we determined how control of expression impacts fitness by measuring exponential growth rates at different expression levels. We reduced β-galactosidase levels by up to 30% using degradation tags, or *lac* operon expression levels by up to 49% by editing the UP element, without affecting growth rates in lactose **(Tables S1** and **S3)**. These results indicate that WT *lac* expression levels are significantly higher than necessary to support maximal growth rates on lactose. Previous work [21] measured expression costs by comparing WT growth rates on glycerol with and without IPTG. However, it was later shown that the cost measured in this way results largely from permease activity in transporting IPTG, and minimal cost was measured in a constitutive, engineered *lac* operon without IPTG, or in the presence of a mutation that disrupts permease function without affecting its expression [25]. Using the native *lac* operon, our results confirm these findings [25] in a wide range of conditions and expression levels **(Fig. S5)**. Using the UP-element library, we detected no difference between *lacI^−^_UE_* and WT*_UE_* growth rates on any carbon source, despite expression levels in *lacI^−^_UE_* increasing up to 6.8 times its levels in lactose **(Table S3)**. We note that at significantly higher expression levels, e.g. when β-galactosidase is over-expressed from a plasmid, expression costs can become substantial [22, 38].

In fluctuating environments, we assessed how sensing impacts fitness by quantifying instantaneous growth rates and lag durations of WT and *lacI^−^*upon metabolic transitions to different carbon sources. Wild-type regulation is susceptible to gene induction lags, which *lacI^−^*avoids by constitutively expressing the *lac* genes **(Fig. 1A)**. However, phenotypic memory in WT cells can attenuate or even eliminate repeated gene induction lags after initial *lac* induction **(Fig. 1A)**, depending on the period between lactose exposures [19]. We showed that selection can act on expression levels to enhance phenotypic memory in fluctuating, but not in constant, environments **(Fig. 1E,F, Figs. S1-S4)** with selective coefficients of ∼1% **(Fig. 1F)**, which declined with increasing duration of environments, consistent with model predictions **(Fig. 1G)**. In addition to avoiding gene induction lags, we found that *lacI^−^* can grow slightly faster than WT upon transition to diauxic growth on lactose and glucose, arabinose, maltose, or xylose **(Fig. 2C-F**, insets). We speculate that this additional advantage of *lacI^−^* stems from its ability to co-utilize both carbon sources, while WT consumes them sequentially [39]. Competition in a fluctuating environment, with periodic glucose and lactose diauxie, showed that *lacI^−^* exhibits a small (∼0.7%) but sustained advantage over WT **(Fig. 3A,B)**, consistent with previous experimental evolution studies that showed that *lac* constitutive mutants rise to fixation in similar conditions [5–7]. We conclude that, to minimize the occurrence of gene induction lags, selection can act to increase phenotypic memory, either by increasing gene expression levels or by losing regulation in favor of constitutive expression.

When *lac* expression levels are above a threshold, we showed that *lacI^−^* cells exhibit a pronounced lag phase upon transition to lactose diauxie **(Fig. 2G-J** and **Fig. S6E-G)**. Such lags were previously reported [26, 27] and later shown to stem from the wasteful flux of lactose through the *lac* permease, which degrades the PMF [28–30]. We termed this behavior ‘catabolite hyperflux lag’ to emphasize its dependence on flux, i.e. a function of both permease expression level/activity and external lactose concentration. To make the connection between catabolite hyperflux lags and selection for regulation, we directly compared *lacI^−^* and WT across multiple conditions and genetic backgrounds. This allowed us to estimate that the threshold *lac* expression level necessary for catabolite hyperflux is ∼3 times the normal expression levels in lactose **(Fig. 2A** and **Table S2)**. By reducing the lactose concentration below the dissociation constant of LacY to lactose (K_D_ = 1 mM [40]), we could reduce hyperflux lag duration, or even eliminate the lag, without compromising growth rate **(Fig. 2J)**. Conversely, by increasing *lac* expression levels via UP element mutations, we could increase hyperflux lag duration by more than 2 hours (see *lacI^−^_UE8_*, **Fig. 4C)**. Importantly, we found that wild-type regulation was not susceptible to catabolite hyperflux lags in any condition **(Fig. 2C-I**, insets), including in genetic backgrounds that increased *lac* expression levels by up to 54% (e.g. WT*_UE8_*, **Fig. 4F)**. This suggests that presence of the repressor is sufficient to maintain *lac* expression levels below the hyperflux threshold across conditions. Competition in a fluctuating environment, with periodic glycerol and lactose diauxie, revealed that WT exhibits a pronounced fitness advantage over the *lacI^−^* strain, which goes extinct within ∼60 generations **(Fig. 3C)**. In light of these findings, our work suggests that periods of glucose starvation, in which bacteria utilize poor carbon sources, combined with transient availability of lactose, are common features of the host niche, which would select for maintenance of the regulator as seen in natural isolates [16].

By sensing and responding to environmental fluctuations, the *lac* repressor enables expression levels to increase gradually, which decreases cAMP levels due to increased lactose catabolism, avoiding over-expression of the permease and concomitant catabolite hyperflux. In this way, the repressor also plays an important role in shaping the environment-dependent fitness landscape in which gene expression levels adapt. In the absence of the repressor, periods of glucose and lactose diauxie select for increasing expression levels while periods of glycerol and lactose diauxie select for decreasing expression **(Fig. 4A,B)**. Such conflicting selection pressures reduce both the efficiency of selection and the maximal long-term growth rate in fluctuating environments. Presence of the repressor enables directional adaptation of gene expression levels across conditions (compare order of curves in **Fig. 4A,B** versus **Fig. 4D,E)**. Additionally, we observed environment-dependent sign epistasis between mutations impacting sensing (WT vs. *lacI^−^*) and control of expression levels (*UE* strains) in the glycerol and lactose diauxic condition **(Fig. 4C,F)**. Due to sign epistasis, as gene expression levels adapt toward higher levels in the presence of the repressor **(Fig. 4F)**, the disadvantage of *lacI^−^* mutations during glycerol and lactose diauxie increases **(Fig. 4C)**, making it increasingly difficult to lose the repressor when such environments recur.

Several questions relevant to our findings constitute key directions for future research. First, how constitutive *lac* cells recover from the catabolite hyperflux lag remains unknown. Previous work showed that the PMF and ATP production can recover after a sufficient period of time [29], and that sodium and potassium gradients play a role in buffering against loss of the proton gradient [30], yet the pathways that underlie recovery have not been identified. By the same token, the inability of certain constitutive mutants to recover from catabolite hyperflux, i.e. cell death or ‘lactose killing’ [27, 28], is not understood mechanistically. To recover from catabolite hyperflux lags, *lacI^−^* cells could metabolize the incoming lactose into glucose, which enables slow growth before the PMF has recovered [41]; while increasing levels of intracellular glucose act to repress LacY activity and decrease cAMP levels thus lowering *lac* expression levels [42]. Interestingly, the presence of non-metabolizable inducers, such as IPTG and TMG, leads to growth rate reduction without causing catabolite hyperflux lags [26, 27, 29]. It has been suggested that this results from a lower intrinsic rate of transport of these inducers by the *lac* permease [30], though this hypothesis remains untested. Lastly, it was shown that in the murine gut, *E. coli* WT and *lacI^−^*strains coexist in lactose diet conditions, while experiments *in vitro* showed that *lacI^−^* had shorter lag times in glucose-to-lactose transitions [43]. It would be interesting to test *in vivo* if in a fluctuating diet between a poor carbon source and lactose the coexistence between the strains is eliminated.

Demand theory of gene regulation argues that constitutive versus regulated expression is selected for based on the overall demand for enzyme expression, which is set by availability of metabolites in the environment [44, 45]. However, the availability of lactose is identical in our competition experiments, and does not determine their outcome **(Fig. 3)**. Instead, we show that the history-dependent physiological state of cells during environmental transitions dictates selection for or against regulation. Our results point to an unexplored role of gene regulation in determining how gene expression levels evolve in fluctuating environments, and suggest that the fitness tradeoffs found here may exist in other gene regulatory networks with similar properties. For example, carbohydrate transport systems in *E. coli* that have a dedicated H^+^ symporter [46] and are positively regulated by Crp (e.g. the *lac, ara, xyl*, *mel*, *dgo*, and *gal* operons) all exhibit negative regulation, suggesting that these systems may be susceptible to catabolite hyperflux. By uncovering the connections between the character of the fluctuating environment, the molecular basis of expression costs, and the role of gene regulation in shaping an evolutionary landscape, our work provides a new basis for understanding and predicting how gene regulatory networks may evolve.

## Acknowledgments

We thank Guillaume Lambert, Keiran Stevenson, and David Specht for sample sequencing and discussions. We thank Yuichi Wakamoto and Reiko Okura for their help with microfluidic device fabrication. T.N. acknowledges support by JSPS KAKENHI Grant Number JP21K20672. This work was supported by the National Institute of General Medical Sciences of the NIH under award numbers R01GM120231 and R01GM148703 to E.K.

## Author contributions

D.P. and E.K. designed research, performed research, analyzed data, and wrote the paper; T.N. analyzed data.

