## Supplementary Information for "Gene regulation, phenotypic memory, and selection in fluctuating environments"

**The PDF file includes:**

Methods
Supplementary Text
Supplementary Figures 1-9
Supplementary Tables 1-4
Supplementary Movies 1,2 Captions

**Other Supplementary Materials for this manuscript include the following:**

Supplementary Movies 1,2
Supplementary Tables 5-7

#### Methods

##### *Escherichia coli* strains and growth conditions

Strain derivatives of *Escherichia coli* MG1655 are listed in **(Table S5)**. Unless stated otherwise, strains were grown in 1X M9 minimal media supplemented to final concentration with 2 mM $MgSO_4$, 100 µM $CaCl_2$ and one of the following carbon source concentrations (w/v) 0.4% lactose, 0.2% glucose, 0.2% glycerol, 0.2% arabinose, 0.2% xylose, 0.4% maltose, 0.4% pyruvate, or 0.4% succinate. In a subset of experiments, strains were grown in M9 minimal media supplemented with 1% casamino acids. Cells were grown with aeration in an incubator set to 37˚C. Whenever applicable, cells were grown with the following antibiotics: carbenicillin 100 µg $mL^{-1}$ or tetracycline 10 µg $mL^{-1}$. When appropriate, 1 mM isopropyl-β-D-1-thiogalactopyranoside (IPTG) was used.

##### Strain construction and plasmids

All chromosomal edits were carried out using Lambda Red Recombineering as previously described [47]. *lacZssrA* strains were previously constructed and described [33]. Briefly, to delete the *lacI* gene, the *tetAsacB* dsDNA selection cassette was first PCR amplified from the T-SACK strain using primers with 35 bp homology flanking the 5' and 3' ends of the *lacI* gene in the chromosome **(see Table S6)**. Colonies were selected after electroporation with the amplified *tetAsacB* cassette on LB plates with 10 µg $mL^{-1}$ tetracycline. Tetracycline positive colonies were tested for toxicity on LB plates supplemented with 6% sucrose. Counter-selection of the *tetAsacB* cassette was performed on LB plates supplemented with 6% sucrose after electroporation of a ssDNA oligonucleotide consisting of the same homology regions that were placed on the primers amplifying the *tetAsacB* cassettes. *lacI*$^-$ genotype of colonies was confirmed using targeted PCR amplification with oligonucleotides **(see Table S6)** outside of the deleted region followed by Sanger sequencing. To generate the modifications in the upstream sequence UP-element region of the *lac* promoter in the WT and *lacI*$^-$ strains, primers with 35 bp homology flanking the 5' and 3' ends of the UP-element region were used to amplify the *tetAsacB* cassette. ssDNA carrying the homologous flanking regions and edited UP elements sequences that previously were shown to alter *lac* expression [36] were used for counter selection as above.

Strains were cured of the pSIM6 Lambda Red plasmid as detailed previously [47]. The MG1655:F3 strain used is identical to the wild-type MG1655 aside from 3 gene deletions (*Δflu, ΔfimA, ΔfliC)* that make the strain suitable for microfluidics experiments. The MG1655:F3 strain and the pPλ-mCherry plasmid were a gift from Yuichi Wakamoto [48]. The T-SACK strain and the pSIM6 plasmids were a gift from Don Court [47]. Plasmids used in this study are listed in **(Table S7)**.

#### Growth dynamics and growth rate measurements

The appropriate strains were taken out on LB plates from -80˚C glycerol stocks and were grown overnight at 37˚C. Each biological replicate corresponds to a single colony of a given *E. coli* strain inoculated in 3 mL of liquid M9 media supplemented with the specified carbon source, as indicated for each condition in figure captions, and grown overnight. Each overnight culture was back diluted 1:100 into the corresponding fresh media and grown to exponential phase. Technical replicates ($n \geq 5$) consist of 8 µL from each of the exponential phase cultures, which were added in a checkerboard pattern to 96-well plates containing 142 µL of the specified media in each well. Four wells on each plate contained only media without bacteria for background measurement. Plates were positioned in a TECAN Spark plate reader (Tecan) and $OD_{600}$ measurements were taken in 5-minute intervals. The plate was shaken continuously inside the plate reader at 37˚C between measurements.

Single growth rate values ($\mu$) were calculated from the slopes of $\ln \mathrm{OD}_{600}$ ranging from 0.01 to 0.1, and doubling time was calculated as $(1/\mu) \ln 2$. Growth rate at time $t$ was calculated as the slope of the $\ln \mathrm{OD}_{600}$ measurements between $t - \Delta t/2$ and $t + \Delta t/2$ using the method of linear least squares, using $\Delta t$ = 180 minutes. Three biological replicates were performed unless otherwise stated in figure captions. For each biological replicate, measurements were averaged over technical replicates on the plate, and the background was subtracted.

For diauxic shift growth measurements, strains were grown as above with a few modifications. Strains were grown overnight in media with a single specified carbon source, and then back diluted 1:100 into fresh media with the same carbon source and grown to exponential phase. Each strain was then inoculated onto wells containing either 1:2 or 2:1 v/v ratio of the specified carbon source to 0.4% lactose. For characterizing catabolite hyperflux lags, *lacI*$^{-}$ cells were grown as above and inoculated into wells containing 1:2 v/v ratio of glycerol and lactose ranging from 0.6% - 0.0125% (11 mM – 0.25 mM). Technical and biological replicates were handled in a similar manner as above. Growth rate and analysis was otherwise identical to the above method.

#### Lag time analysis

WT and *lacZssrA* strains were plated on LB plates from -80˚C glycerol stocks and grown at 37˚C overnight. Single colonies were inoculated in 3 mL of liquid M9 minimal media with 0.2% w/v glucose and were grown with aeration overnight. Cells were diluted 1:100 into 30 mL of M9 media with 0.2% glucose and grown until cultures reached an $OD_{600}$ of 0.1. Cells were then spun down in 50 mL tubes at 6,000 x g in a tabletop centrifuge for 10 minutes to remove the supernatant containing M9 glucose media. Cells were washed twice in M9 media lacking any carbon source and then resuspended in 400 µL (75X concentration) of M9 media without any carbon source. Technical replicates (n ≥ 12) consist of 2 µL from each of the exponential phase cultures, which were added in a checkerboard pattern to 96-well plates containing 148 µL of the specified media in each well. Two wells on each plate contained only media without bacteria for background measurement. Plates were positioned in a TECAN Spark plate reader (Tecan) and $OD_{600}$ measurements were taken in 2-minute intervals. The plate was shaken inside the plate reader at 37˚C continuously between measurements. Lag time was calculated by fitting the $OD_{600}$ (< 0.2) with $OD = A\ (\text{for}\ t \leq T)\ ,\ Ae^{r(t-T)}\ (\text{for}\ t > T)$, where $A$ = initial $OD_{600}$, $r$ = growth rate, and $T$ = lag time, using nonlinear least squares fitting. The analysis is conducted using Julia with the LsqFit.jl package.

#### Competition experiments and strain frequency analysis using barcode sequencing

*lacZssrA* strains were plated on LB plates from -80˚C glycerol stocks and grown at 37˚C overnight. Single colonies from each strain were grown overnight in M9 media supplemented with either 0.2% glucose or 0.4% lactose w/v. Each strain was normalized to an $OD_{600}$ of 1, and 1 mL of each of *lacZssrA* strains from either the glucose or lactose media were separately combined and vortexed. Separate tubes were mixed from the strains grown in glucose for the G/L fluctuation condition. 1 mL of the combined strains was sampled from each condition into tubes and the supernatant was removed by centrifugation (15,000 x g, 2 minutes) and cell pellets were placed in -20˚C for storage. These samples mark day 1 of the batch competition experiment. Cells were grown in one of three conditions: constant glucose (G), constant lactose (L), or daily fluctuations between glucose and lactose (G/L). Three parallel series of competition experiments were run using dilutions 1:16, 1:100, and 1:10,000. Each day at the same time $OD_{600}$ was measured and 1 mL of cells were sampled from each culture into tubes, supernatant was removed after centrifugation (15,000 x g, 2 minutes) and the samples were placed at -20˚C for storage. $OD_{600}$ measurements of each culture were recorded, and cells were diluted appropriately according to the specific condition in each experiment. DNA was extracted from each sample using Wizard Genomic DNA Purification Kit (Promega) according to manufacturer instructions. Purified DNA was normalized to 100 ng/µL using double-distilled $H_2O$ and amplified by PCR using Platinum

SuperFi II DNA Polymerase-High Fidelity PCR enzyme according to manufacturer instructions (Thermo Fisher Scientific). A combination of ssDNA oligos was used, which contained unique identifiers for each cell dilution, growth condition, and day **(see Table S5)**. In addition, TruSeq UDI indexes were added for downstream sequencing. Each PCR amplified sample was verified on a gel for the correct length. Prepared libraries were diluted to 50 pM and sequenced on an Illumina iSeq 100 platform with 2 x 150 bp paired-end reads. Raw reads were analyzed by trimming Illumina adapters and aligned to each *lacZssrA* variant sequence. Frequency of each *lacZssrA* strain was calculated by dividing each *lacZssrA* read count by the total *lacZssrA* reads. The strain frequency analysis was implemented in Julia with FASTX.jl and BioSequences.jl packages.

#### Fitting competitive dynamics using a relative exponential growth model

Strain frequency dynamics were fit assuming that each strain grows exponentially with a rate determined by its relative selection coefficient, denoted by $s_i$, where $i$ indexes the strain. We let $x_i(t)$ denote the predicted frequency of strain $i$ at time $t$, where $\sum_i x_i(t) = 1$, the summation running over all strains in the competition ($i = 1, 2, \ldots, 6$). Assuming independent exponential growth of each strain, where $c_i$ is the frequency of strain $i$ at $t = 0$, we obtain

$$x_i(t) = \frac{c_i e^{s_i t}}{\sum_j c_j e^{s_j t}} .$$

The free parameters in the model are $c_i$ and $s_i$, for $i = 1, 2, \ldots, 6$, where $s_i$ are determined up to an additive constant. We denote by $y_i(t)$ the experimentally-measured frequency dynamics of each strain. Non-linear least squares fitting was performed to minimize the squared deviation $\sum_t \sum_i [y_i(t) - x_i(t)]^2$ over the set of parameters using the quasi-Newton method implemented in Wolfram 14.1. Because each trajectory $y_i(t)$ is determined by two parameters ($c_i$ and $s_i$) together with the overall normalization constraint, fitting converges rapidly. Since the time unit for $t$ is days in each experiment, we initially infer values of $s_i$ having units of 1/d. The number of generations (gen) per day is calculated from the dilution factor in each experiment: 1:16 = 4 gen/d, 1:100 = 6.6 gen/d, 1:10,000 = 13.3 gen/d. We use these conversion factors to report $s_i$ in units of 1/gen for each experiment. Because values of $s_i$ are determined only up to an additive constant, in each experiment we calculate the mean inferred selection coefficient over the six strains, $\bar{s} = (1/6)\sum_i s_i$. We then report selection coefficients relative to the mean in the experiment, i.e. $s_i - \bar{s}$.

#### Whole-genome sequencing and variant calling for isolates from competition experiments

Whole-genome sequencing and variant calling were used to determine the mutational landscape of selected isolates from the phenotypic memory selection experiments. Individual colonies were isolated from day 14 competition samples and DNA was extracted as above. For reference comparison, whole-genomes of the respective parental strains (*AAV, NYNY,* or *No Deg*) were prepared as well. DNA fragmentation and library preparation were performed on individual whole-genomes according to Illumina guidelines. Libraries were sequenced on the Illumina Novaseq X platform achieving a mean genome coverage of over 150x. FASTQ files were generated and trimmed sequences were aligned to the *E. coli* reference genome GCF_000005845.2 (RefSeq NC_000913.3). Sentieon DNASeq variant calling was used to determine single nucleotide polymorphism (SNPs) and structural variants including indels [49]. Summary of SNPs and structural variants among the isolates is shown in Table S4.

#### Modeling *lacZssrA* strain competitions in a periodic environment

A biophysical model of growth and Lac protein expression in lactose was previously developed for the *lacZssrA* strains [33]. Briefly, the ordinary differential equation (ODE) model uses LacZ protein levels as a proxy for cells' metabolic capacity, which is represented by a Monod function of LacZ concentration. It models the dynamics of LacZ protein expression taking into account a changing production rate, an ssrA tag-dependent degradation rate, and a growth-rate-dependent dilution rate. Model parameters were inferred from LacZ induction kinetics of each of the *lacZssrA* strains [33]. Here, we used the same parameters to predict selection coefficients of the *lacZssrA* strains in a glucose-lactose fluctuating environment **(Fig. 1G)**. In lactose environments, the changing population growth rate, $\Lambda(t)$, was modeled as $\Lambda(t) = \Lambda_{max} Z(t)^h/(K^h + Z(t)^h)$, where $\Lambda_{max}$ is the maximal growth rate in lactose, $Z(t)$ is the changing LacZ expression level in Miller units, $K$ is the level of LacZ that supports half-maximal growth in lactose, and $h$ is the Hill coefficient ($h = 1$ was used [33]). In glucose environments, the growth rate $\Lambda(t)$ was set to a constant value determined by the measured doubling time in glucose; and LacZ production rate was set to 0, hence $Z(t)$ decayed due to degradation and dilution. We numerically solved the ODE model in a periodic environment, where the duration $\tau$ of each environment was given by the number of generations per day ($n_{gen} \equiv \log_2 D$, where $D$ is the dilution factor) times a mean doubling time of 66 min. We ran the model until the long-term dynamic steady-state is reached, and then averaged the growth rate over a single period to predict the selection coefficient of the $i$-th strain, $s_i = \frac{1}{2n_{gen}} \int_t^{t+2\tau} \Lambda(t)dt$. This yields the selection coefficients in units of 1/gen. To determine the Hill coefficient, which impacts growth lags, we compared predicted values of $s_i$ with values measured from competition experiments **(Fig. 1G)** for different values of $h$. We found that the original model ($h = 1$) yielded the same trend shown in Fig. 1G, but systematically underestimated the relative selection coefficients by ~0.3%; the best overall fit was obtained with $h = 3$, which is shown in Fig. 1G.

To predict the population dynamics in the competition experiments (shown in **Fig. S4C)**, we simultaneously solved the ODE models for all six strains in a mixed culture. We introduced the competitive interaction between strains using a single shared resource (glucose or lactose) which is consumed by each strain proportionally to its frequency and growth rate. As the resource is globally consumed, the growth rates of each strain decay as a Monod function of the shared resource; the initial concentration of the resource thus determines the number of generations before saturation. The initial frequencies at $t = 1$ for each strain were chosen to match the experimentally measured values from Fig. 1E, and the predicted frequencies ($t > 1$) are shown as points in Fig. S4C. We fit the relative exponential growth model to the simulated frequencies to infer the relative selection coefficients shown in Fig. S4G.

#### Microfluidic device fabrication

Using standard photolithography techniques, we fabricated a polydimethylsiloxane (PDMS) device similar to those used in [18, 19], with some modifications. The device consists of a T-shaped main flow channel and 1600 growth channels. The approximate dimensions of the main channel and the growth channel are 16 mm (L) × 200 μm (W) × 20 μm (H) and 65 μm (L) × 7.5 μm (W) × 1.3 μm (H), respectively. A circular inlet/outlet (ϕ=500 μm) is located at each end of the main channel, and two inlets are spaced approximately 7 mm apart. Growth channels are approximately 7.5 μm apart.

We created the CAD designs of the main channel and the growth channels using KLayout. These designs were printed on photoresist-coated chrome-on-glass masks (CBL4006Du-AZP, Clean Surface Technology) using a maskless aligner (μMLA, Heidelberg). The UV-exposed regions of the resist (AZP1350) were removed by NMD-3 (Tokyo Ohka Kogyo), and the exposed chromium was etched by MPM-E350 (DNP Fine Chemicals). After removing the remaining photoresist layer using acetone (Wako), the masks were rinsed with MilliQ water and air-dried.

An SU-8 mold for the PDMS device was fabricated on a silicon wafer (76.2 mm, ID 447, University Wafer) in two steps. For the first layer or growth channels, the wafer was spin-coated with SU-8 (Kayaku Advanced Materials) using a spin coater (MS-A150, Mikasa) at 500 rpm for 10 seconds, ramped to 1400 rpm over 90 seconds, and then at 1400 rpm for 30 seconds. The SU-8-coated wafer was baked at 65°C for 1 min and then at 95°C for 3 min. The SU-8 layer was exposed to

UV light three times every 10 sec (22.4 mW/cm2 for 1.7 sec per exposure) using a mask aligner (MA-20, Mikasa). The post-exposure wafer was baked at 65 °C for 1 min and then at 95 °C for 2 min. The wafer was developed with SU-8 developer (Kayaku Advanced Materials), rinsed with 2-propanol (Wako), and air-dried. The height of the growth channel was measured as 1.27 using a stylus profiler (DektakXT-A, Bruker). For the second layer, the same wafer was spin-coated with SU-8 3025 (Kayaku Advanced Materials) at 500 rpm for 10 seconds and then at 3,000 rpm for 30 seconds. The wafer was baked at 65°C for 1 min and then at 95°C for 7 min. The second SU-8 layer was exposed to UV light (22.4 mW/cm2 for 20 sec). The post-exposure wafer was baked at 65°C for 1 min and then at 95°C for 7 min. The wafer was developed with SU-8 developer, rinsed with 2-propanol, and air-dried.

Lastly, we created an epoxy-based replica of the SU-8 master, adapting the Jun lab's protocol for making an epoxy mold [https://github.com/junlabucsd/mother-machine-protocols]. The PDMS base and curing agent (Sylgard 184 Silicone Elastomer Kit, Dow) were mixed at a 10:1 ratio, poured onto the SU-8 mold in a petri dish, and degassed for 1 hour using a vacuum desiccator. The PDMS plate was cured at 65°C overnight. At the same time, 10 g of PDMS mixed at a 20:1 ratio was poured into the bottom half of a new petri dish, and it followed the same degassing and curing procedures. This PDMS pad served as the base on which the devices would sit, and the epoxy was poured. The device was peeled off the mold and washed in ethanol using sonication for 30 min, followed by drying at 65°C. The hardened PDMS pad and the device were cleaned using Scotch tape. The device was placed on the PDMS pad with the feature side up. The epoxy (Loctite E-30CL Hysol epoxy) was mixed using a dispenser gun and a static mixing tube in a plastic cup. The epoxy was degassed for 10 minutes and poured over the PDMS device, which was sitting on a PDMS pad in the petri dish. After sitting overnight, the petri dish, PDMS device, and PDMS pad were removed to get the epoxy duplicate of the SU-8 master.

#### Time-lapse microscopy with microfluidic device

A PDMS device was peeled off the epoxy duplicate described above. Media input and output holes were punched into the designated areas of the PDMS device and then attached via oxygen plasma to a 50 x 7mm glass bottom petri dish. The WT and *lacI*⁻ strains were transformed with pPλ-mCherry to allow fluorescence tracking during growth. Each strain was initially grown in 3 mL of M9 media supplemented with 0.2% glucose and 1% casamino acids from overnight culture. At a cell density of $OD_{600}$ = 0.3, 1 mL of cells were pelleted by centrifugation (15,000 x g for 2 minutes) and resuspended with 100 µL of M9 glucose media supplemented with 1% casamino acids. The concentrated cells were then injected into the PDMS device and incubated at 37˚C until all chambers were filled (~1 h). Tygon tubing was attached to syringe needles and to each hole of the PDMS device, and connected to solenoid valves. Pressurized media (M9 supplemented with 1% casamino acids and 0.2% glucose or 0.4% lactose) was pushed through the main channel at a flow rate of 5 mL $h^{-1}$ and 6 PSI. The connected device was placed within a temperature-controlled enclosure on a stage of an inverted Nikon Eclipse Ti2 microscope equipped with a Plan Apo 20x NA 0.75 objective and DSQi2 camera, and images were acquired using the NIS Elements 5.10.01 software. Automatic media change was carried out using a flow control manager script implemented in µManager [50]. Each strain was initially grown in glucose media and fluctuations between glucose and lactose were performed in 4.5-hour intervals. Cells were tracked by fluorescence excitation of 555 nm and 629.5 nm emission and images were taken every 2 minutes from each chamber.

#### Evaluating growth rate from time lapse images

Growth rates were calculated as the slope of the optical flow of fluorescence images along the direction of cell elongation (vy) vs distance from the closed end of a chamber (y), as previously done [18]. Using a custom ImageJ script, after the original fluorescence images were stabilized to maintain the location of each growth channel across different time frames, the rectangle surrounding a cell population was extracted for every growth channel. For each rectangle at each time point, frame-to-frame velocity vector fields were computed using Farnebäck's optical flow algorithm [51], which is implemented in Julia with Images.jl package. In each rectangular region, vy was averaged in the direction parallel to the main channel and further averaged over a 10-minute time window. Then, their mean and standard deviation were calculated over all chambers. Finally, the mean growth rate and its standard error are computed at each time point using weighted least squares in 10 μm < y < 30 μm. The growth rate calculations following optical flow analysis were implemented in Julia with LsqFit.jl package.

#### β-Galactosidase activity measurements using the Miller assay

Modified Miller assay was performed as previously described [52, 53]. Briefly, after $OD_{600}$ measurements, 40 μL of cells were placed in a tube containing 60 μL of permeabilization solution consisting of 100 mM of $Na_2HPO_4$, 20 mM KCl, 2 mM $MgSO_4$, 0.8 mg $mL^{-1}$ CTAB (hexadecyltrimethylammonium bromide), 0.4 mg $mL^{-1}$ sodium deoxycholate, and 5.4 μL $mL^{-1}$ β-mercaptoethanol. At the end of sample collection, 600 μL of substrate solution containing 60 mM $Na_2HPO_4$, 40 mM $NaH_2PO_4$, 1 mg $mL^{-1}$ o-nitrophenyl-β-D-galactopyranoside (ONPG) and 2.7 μL $mL^{-1}$ β-mercaptoethanol were added to each tube and samples were incubated for 10 minutes at room temperature. After 10 minutes, 700 μL of stop solution containing 1 M $Na_2CO_3$ were added to each sample. Samples were then centrifuged at 15,000 x g in a tabletop centrifuge for 5 minutes and 1 mL of supernatant from each sample was placed in cuvettes and measured at $OD_{420}$ using a GENESYS 140/150 spectrophotometer (Thermo Fisher Scientific). β-Galactosidase activity measurement is given by, $1000 \cdot \frac{Abs_{420}}{Abs_{600} \cdot v \cdot t}$, $t$ = 10 min, and v = 0.04 mL for all samples; and its unit is known as Miller units (MU). All Miller measurements were performed in biological triplicates.

#### Oxygen consumption rate measurements

WT and *lacI*$^{-}$ strains were plated on LB plates from -80˚C glycerol stocks and grown at 37˚C overnight. A night prior to each experiment, Seahorse Sensor Cartridge plate was loaded with 200 µL of calibrant solution to hydrate the sensors according to manufacturer instructions. Single colonies from each strain were grown overnight in M9 media supplemented with either 0.2% glucose or 0.2% glycerol w/v. Each overnight culture was back diluted 1:100 into the corresponding fresh media and grown for an hour before adding 1 mM IPTG to a subset of the cultures. Cultures were then incubated at 37˚C and grown to exponential phase. $OD_{600}$ measurements were taken, and cultures were back diluted with the appropriate media to $OD_{600}$ = 0.01. 90 µL of each sample were added to a Poly-D-Lysine (PDL) culture microplate. Rows A and H were filled with only media and cells were added in 6 technical replicates into the rest of the wells in a checkboard pattern. To adhere cells to the bottom of each well, plates containing cells were centrifuged at 1,400 x g for 10 minutes using a tabletop centrifuge with low deceleration setting to avoid disruption of the attached cells. 90 µL of the appropriate media were then added for a total volume of 180 µL. Hydrated Sensor Cartridge plate was loaded onto a Seahorse XF Pro Flux Analyzer (Agilent) according to manufacturer guidelines and the plate containing cells was loaded afterwards. After initial equilibration, measurements were acquired over 3 cycles (2.5 minutes mixing and 2.5 minutes measurements) for a total of 15 minutes. Seahorse measurements were performed in biological triplicates. Each experiment was performed on a different day with independent bacterial cultures.

#### Colony forming unit (CFU) measurements

To asses if lactose-induced cell death occurred when plating the *lac* constitutive strain on lactose media as previously found [28], we compared the colony forming unit (CFU) counts of our *lacI*$^{-}$ and WT strains. WT and *lacI*$^{-}$ were plated on LB plates from -80˚C glycerol stocks and grown at

37˚C overnight. Single colonies from each strain were grown overnight in M9 media supplemented with 0.2% glycerol w/v. Each overnight culture was back diluted 1:100 into M9 media supplemented with 0.2% glycerol and grown to exponential phase. 100 µL were taken for plating after serial dilution in M9 media without any carbon source. Cells were plated on either LB or M9 0.4% lactose media and incubated overnight. Individual colonies were counted, and CFU/ml was calculated. Each biological replicate (n = 3) consists of (n = 3) technical replicate plates.

To measure CFUs during catabolite hyperflux lag and recovery phase, the *lacI*$^-$ strain was grown as above. After reaching exponential growth in M9 media supplemented with 0.2% glycerol, at $t$ = 0, cultures were diluted to $OD_{600}$ = 0.01 in M9 supplemented with 0.4% lactose. Samples were taken every 30 minutes during the catabolite hyperflux lag (0 – 120 minutes) or every hour during the growth recovery phase (120 – 240 minutes). For each sample, $OD_{600}$ measurements were taken and CFU counts were performed as above.

#### Diauxic shift competitions

The WT, *NYNY*, and *lacI*$^-$ strains were plated on LB plates from -80˚C glycerol stocks and grown at 37˚C overnight. Single colonies from each strain were grown overnight in M9 media supplemented with either 0.2% glucose or 0.2% glycerol w/v. Each strain was normalized to an $OD_{600}$ of 1, and 1 mL of each strain from either the glucose or glycerol media were separately combined and vortexed. A separate tube was mixed from the strains grown in glucose or glycerol for the glucose-to-glucose/lactose diauxie or the glycerol-to-glycerol/lactose diauxie fluctuations, respectively. These samples mark day 1 of the batch competition experiments. Cells were grown in 1:100 dilution in 4 conditions: (i) constant glucose, (ii) constant glycerol, (iii) daily fluctuations between glucose and glucose/lactose diauxie, and (iv) daily fluctuations between glycerol and glycerol/lactose diauxie. Each day at the same time $OD_{600}$ was measured and 30 µL were added to tubes with 3 mL fresh media. Every other day, after all cultures were grown in a single carbon source overnight, 100 μL were taken for plating after serial dilution in M9 media without any carbon source. Cells were plated on M9 minimal media supplemented with 0.2% glucose and 2 μg $mL^{-1}$ 5-bromo-4-chloro-3-indolyl-β-D-galactopyranoside (X-gal), which is an analog of lactose that can be hydrolyzed by β-galactosidase to yield blue colonies [52, 54]. The X-gal indicator plates allowed us to distinguish between the *lacI*$^{-}$ strain (constitutive *lac* operon expression) and the WT and *NYNY* strains (negatively regulated). For each growth condition, cells were plated onto three plates serving as technical replicates. Each plate was then placed for overnight growth at 37˚C. All the blue and white colonies were counted and a randomly chosen subset (n ≥ 8) of the white colonies from each plate was analyzed by PCR to distinguish between the WT and *NYNY* strains. Specifically, the 3' end of the *lacZ* gene was amplified by PCR using oligonucleotides **(see Table S2)** and PCR products were run on a 2% agarose gel. These analyses were repeated for the duration of the experiment (14 days). Strain frequency was calculated based on the X-gal and PCR colony counts.

#### Statistical analysis

When appropriate, statistical analysis was performed using Prism 10.5 (GraphPad Software) and Wolfram 14.1 as indicated in figure captions.

### Supplementary Text

#### Supplemental experiments for selection of phenotypic memory

A pilot competition experiment was run for 5 days at a single daily dilution (1:100) in three different conditions: constant glucose, constant lactose, or daily alternation between glucose and lactose (G/L); sampling was performed twice daily **(Fig. S1)**. Selection against lower *lac* expression levels was observed in the G/L condition **(Fig. S1F)**, while no selective differences were measured between strains in either constant glucose **(Fig. S1D)** or constant lactose **(Fig. S1E)**.

Longer competition experiments were run for 14 days at three different daily cell dilutions (1:16, 1:100, 1:10,000) in each of the three media conditions. The strength of dilution determines the number of generations (gen) per day: 1:16 = 4 gen, 1:100 = 6.6 gen, and 1:10,000 = 13.3 gen. In the alternating condition, the number of generations corresponds to the effective duration of each environment. Since *lac* operon phenotypic memory is lost only by protein dilution due to cell division, the number of generations spent in the absence of *lac* induction (i.e. in the glucose environment) directly impacts memory levels [19]. We previously showed that cells avoid lag phases in the alternating condition when glucose environments last up to 4 generations [19]. Lag phases are observed when glucose exposure lasts >4 gen, and increase in length with increasing number of generations spent in glucose. After >10 gen, cells exhibit a full lag phase at each lactose encounter. Reducing β-galactosidase expression levels is therefore expected to have a more pronounced effect on the lag phase for shorter glucose exposures.

Due to the larger number of generations in the 14 day experiments relative to the pilot, there is a higher likelihood that beneficial mutations unrelated to *lac* operon expression could rise in frequency during these experiments. This effect was apparent in all three of the 1:10,000 dilution experiments, which had the largest total number of generations. In each of the 1:10,000 runs **(Fig. S2C, S3C,** and **S4B)**, we observed a different *lacZssrA* strain rising in frequency in the second half of the experiment, and reaching a frequency of >80% by day 14. To determine which mutations increased in these experiments, we isolated three colonies of the dominant *lacZssrA* strain from each of the respective day 14 samples, and performed whole-genome sequencing followed by variant calling. SNPs and structural variants are summarized in Table S4, and no mutations were found in the *lac* operon.

Similar behavior was observed in one of the 1:100 experiments, in which the *No Deg* strain rapidly increased in frequency after day 10, reaching a frequency of 0.33 at day 14 **(Fig. S3B)**. We initially performed whole population sequencing on genomic DNA extracted from the day 14 time point. We searched for SNPs and/or structural variants including insertions and deletions that were enriched to a frequency ~0.3, similar to the *No Deg* strain's frequency at that time point, but did not find such variants. We then isolated six *No Deg* colonies from the same samples, performed whole-genome sequencing, and mapped SNPs and structural variants. We found that compared to the ancestor (day 1, *No Deg* strain), there were two SNPs and one structural variant that were shared across multiple isolates **(Table S4)**. We found an IS1F mobile element insertion in the *recG* gene in 3/6 the colonies. To rule out potential *lac* operon-specific mutations, we isolated an additional nine *No Deg* colonies and performed targeted Sanger sequencing on the entire *lac* operon. We found no mutations compared to the *No Deg* reference or MG1655 parental strain, aside from the *ssrA* tag that we had inserted at the 3' end of the *lacZ* gene.

#### Doubling time in IPTG-supplemented arabinose media

We found that the addition of IPTG to the WT and *lacI*$^-$ strains growing in 0.2% arabinose resulted in a substantial increase in doubling time, which was significantly higher than in any other carbon source we tested **(Fig. S5B)**. It was previously shown that lactose and the synthetic *lac* operon inducer TMG (methyl-β-thiogalactoside) can enter *E. coli* cells through the arabinose transport systems [55]. We hypothesize that IPTG can enter through either the ATP dependent high-affinity arabinose transport complex (AraFGH), the low-affinity arabinose-proton symporter (AraE), or both. This could result in a synergistic reduction in growth rate due to IPTG transport by both LacY and arabinose transport systems, leading to PMF and ATP depletion.

#### Supplementary Figures

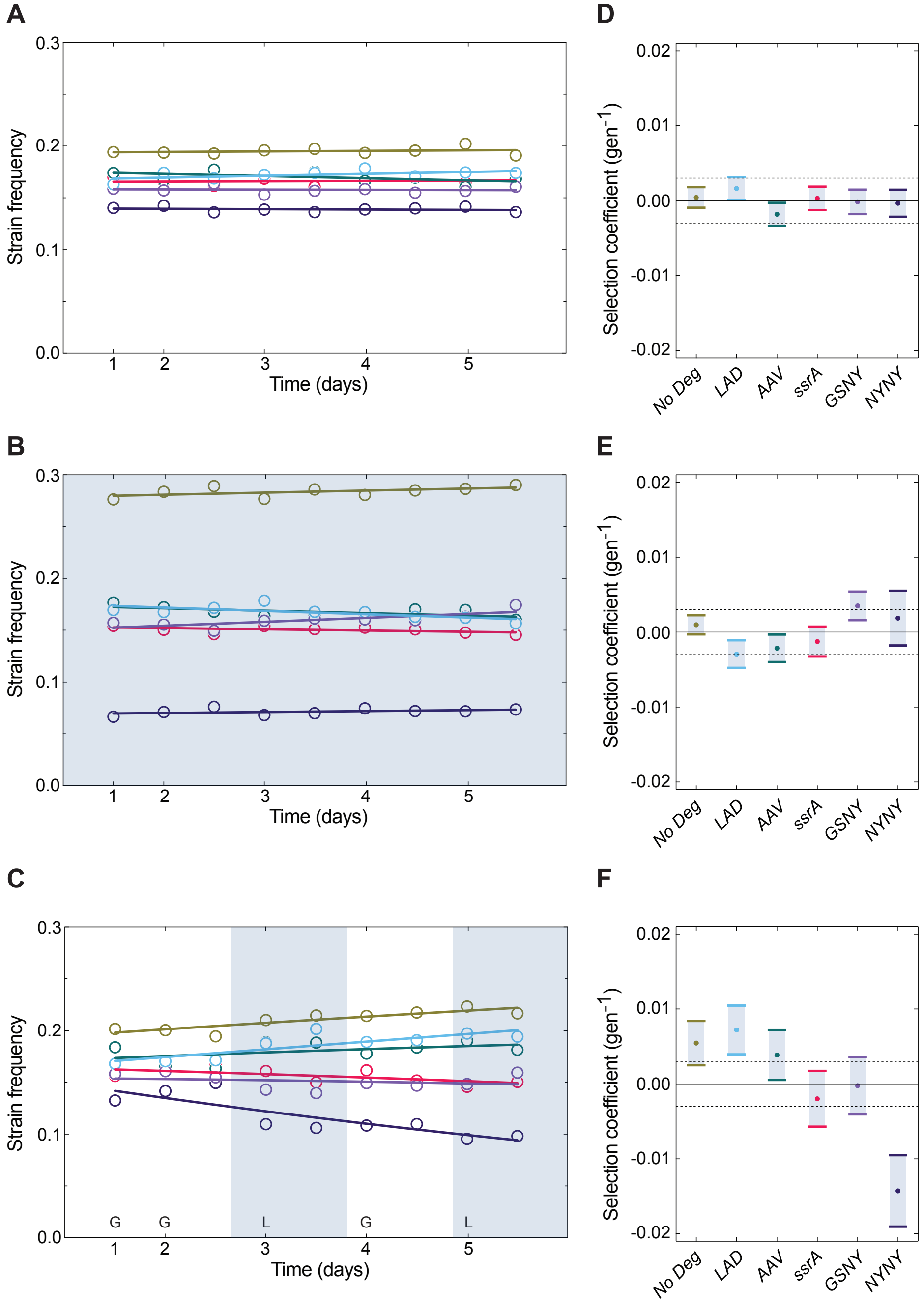


**Fig. S1. Pilot competition experiments for each condition at a single dilution.**
Results from a 5-day pilot experiment using a 1:100 daily dilution factor are shown. Samples were taken twice a day, once right before daily dilution (stationary phase) and once at 5 h post dilution (exponential phase). Competition experiments were run in **(A)** glucose, **(B)** lactose, or **(C)** daily alternation between glucose or lactose (G/L). A relative exponential growth model of the competition was fit (solid lines) to infer selection coefficients. **(D-F)** Relative selection coefficients (points) and 95% confidence intervals (bars) are shown for each strain inferred from model fitting in panels A-C respectively. Dashed line indicates an upper and lower bound of confidence intervals inferred from competition experiments in constant glucose of the pilot experiments (Fig. S1D).

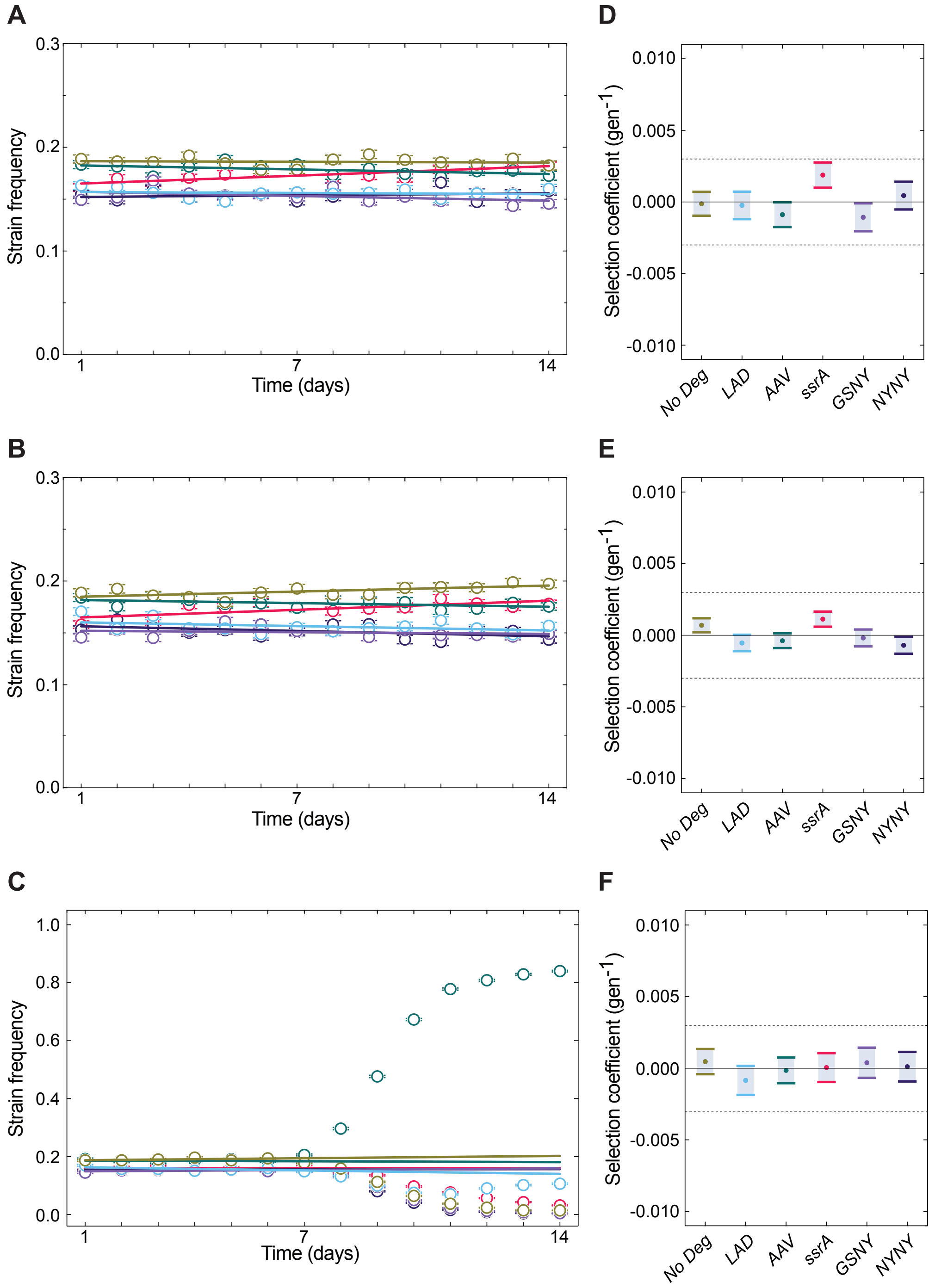


**Fig. S2. Competition experiments in constant glucose at three different dilutions.**
Strain frequency of each *lacZssrA* strain in glucose in either 1:16 **(A)**, 1:100 **(B)**, or 1:10,000 **(C)** daily dilution is shown over two weeks. A relative exponential growth model of the competition was fit (solid lines) to infer selection coefficients. **(D-F)** Relative selection coefficients (points) and 95% confidence intervals (bars) are shown for each strain inferred from model fitting in panels A-C respectively. Due to the rise of mutations in the 1:10,000 experiment at day 7, fitting in panel C and selection coefficients in panel F only use data from days 1 – 6. Dashed line indicates an upper and lower bound of confidence intervals inferred from competition experiments in constant glucose of the pilot experiments (Fig. S1D).

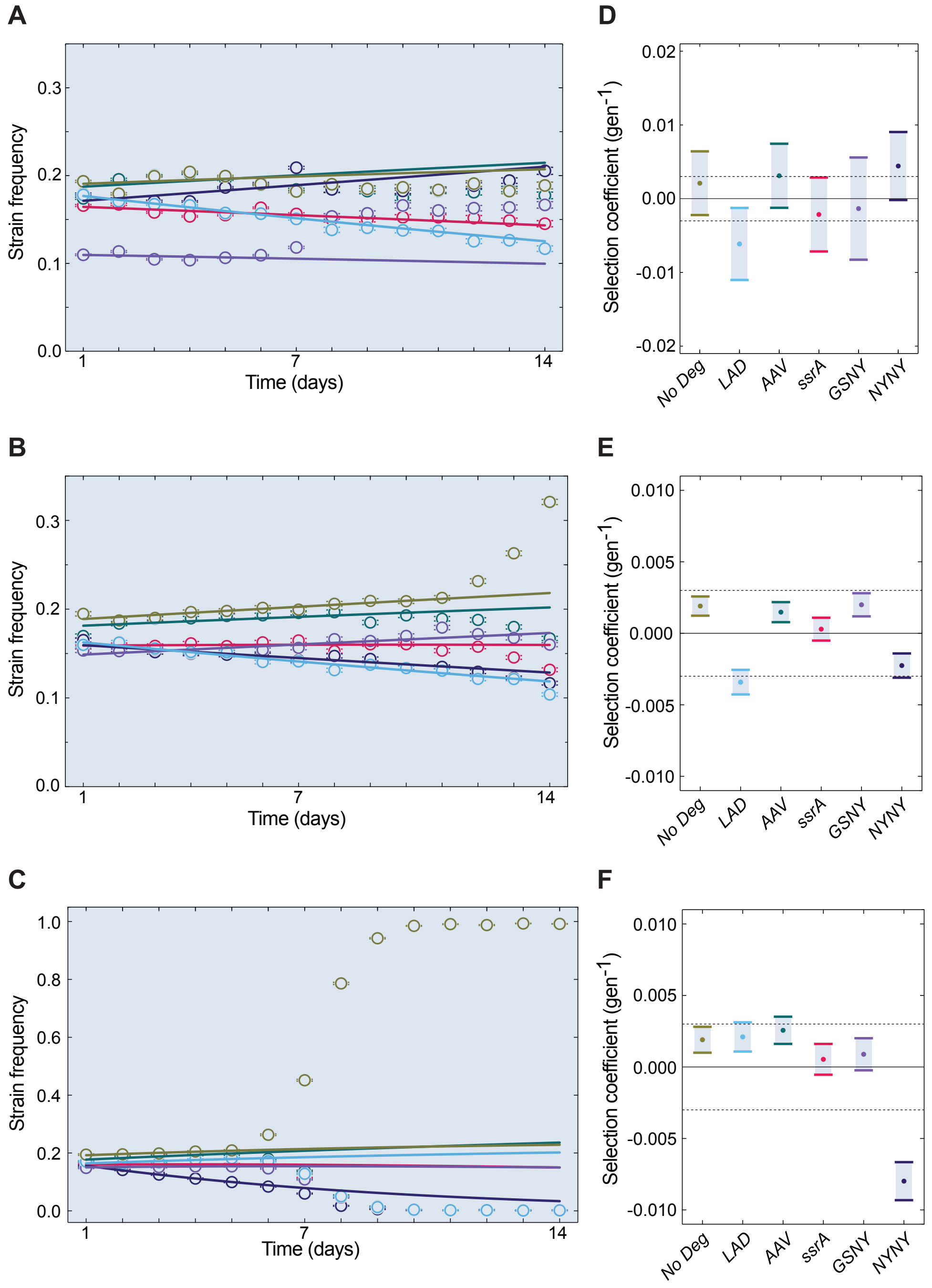


**Fig. S3. Competition experiments in constant lactose at three different dilutions.**
Strain frequency of each *lacZssrA* strain in lactose in either 1:16 **(A)**, 1:100 **(B)**, or 1:10,000 **(C)** daily dilution is shown over two weeks. A relative exponential growth model of the competition was fit (solid lines) to infer selection coefficients. **(D-F)** Relative selection coefficients (points) and 95% confidence intervals (bars) are shown for each strain inferred from model fitting in panels A-C respectively. Due to the rise of mutations in the second half of the experiments, fitting for the 1:16, 1:100, and 1:10,000 experiments only use data from days 1 – 6, 1 – 10, and 1 – 5, respectively. Dashed line indicates an upper and lower bound of confidence intervals inferred from competition experiments in constant glucose of the pilot experiments (Fig. S1D).

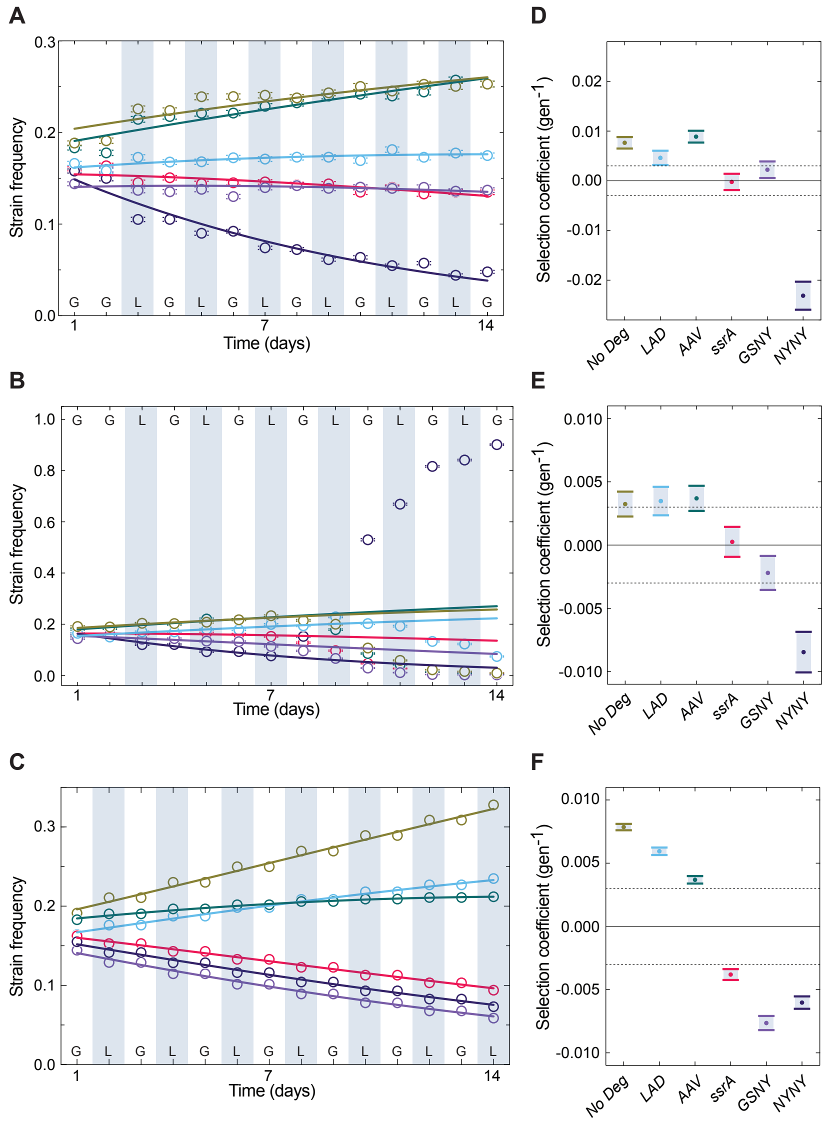


**Fig. S4. Competition experiments in a fluctuating environment at additional dilutions and model prediction.**

Strain frequency of each *lacZssrA* strain in alternating glucose/lactose (G/L) in 1:16 **(A)** or 1:10,000 **(B)** daily dilution is shown over two weeks. A relative exponential growth model of the competition was fit (solid lines) to infer selection coefficients. **(D,E)** Relative selection coefficients (points) and 95% confidence intervals (bars) are shown for each strain inferred from model fitting in panels A,B. Due to the rise of mutations in the 1:10,000 experiment at day 8, fitting in panel B and selection coefficients in panel D only use data from days 1 – 7. Dashed line indicates an upper and lower bound of confidence intervals inferred from competition experiments in constant glucose of the pilot experiments (Fig. S1D). **(C)** Using a biophysical model we previously developed [33], we predicted the competitive dynamics of the strain library over the 14-day experiment. The initial frequency of each strain was set to match the measured frequencies at Day 1 in the fluctuating G/L competition experiment with 1:100 dilution (shown in Fig. 1E). Each point corresponds to the predicted frequency at the beginning of each day. We fit the relative exponential growth model (solid lines) to the model's predicted frequencies (points) to infer selection coefficients. **(F)** Relative selection coefficients (points) and 95% confidence intervals (bars) are shown for each strain inferred from the fit shown in panel C.

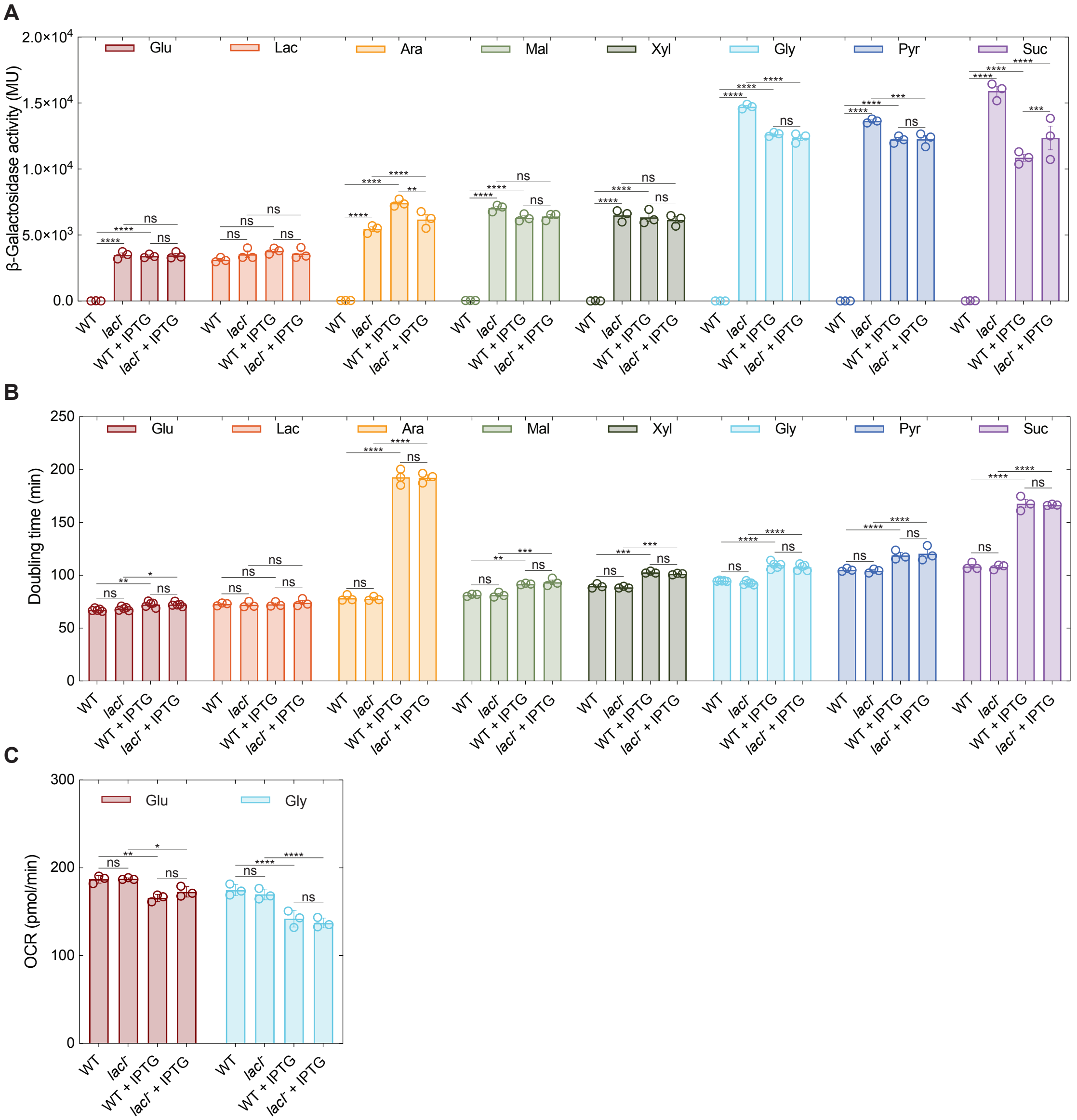


**Fig. S5. *lac* operon expression, growth measurements, and bioenergetics of WT and *lacI*$^-$ strain in various growth conditions.**

**(A)** β-Galactosidase activity levels in Miller units (MU) are shown for WT and *lacI*$^-$ strains grown in minimal media supplemented with either glucose, lactose, arabinose, maltose, xylose, glycerol, pyruvate, or succinate and with or without 1 mM IPTG. Bars indicate mean ± SD (n = 3). Data without IPTG is also shown in Fig. 2A. **(B)** Doubling time measured for WT and *lacI*$^-$ strains in the same conditions as panel A. Individual biological replicates are shown in circles (n ≥ 3). See additional discussion of these results in supplementary text. Data without IPTG is also shown in Fig. 2B. **(C)** Oxygen consumption rate (OCR, in picomoles of molecular oxygen per minute) is shown for WT and *lacI*$^-$ cultures grown in minimal media supplemented with glucose or glycerol, with or without IPTG. Measurements were acquired using a Seahorse XF Pro Flux Analyzer. OCR data is shown for averages of $t$ = 0 – 10 minutes ± SD (n = 3). Two-way ANOVA with Tukey multiple comparisons test was used for statistical analysis. Statistically significant differences are indicated ****$p$<1e-4, ***$p$<1e-3, **$p$<1e-2, *$p$<0.05, ns = $p$>0.05.

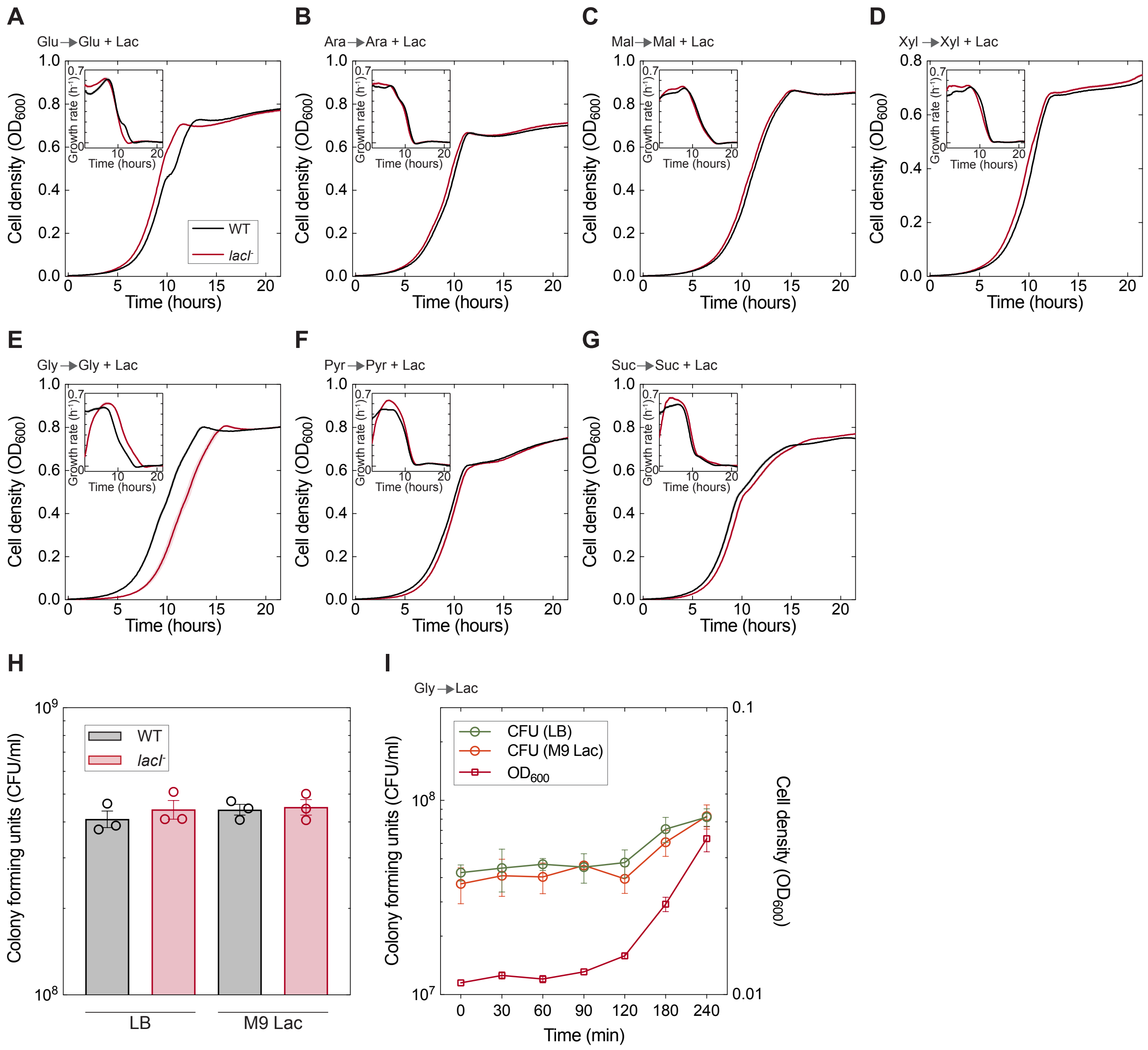


**Fig. S6. Additional diauxic growth dynamics and CFU counts related to catabolite hyperflux lags.**

Diauxic growth dynamics of WT and *lacI*$^-$ strains. $OD_{600}$ **(A-G)** and instantaneous growth rates **(insets)** shown as mean (solid line) ± SEM (shaded envelope) over biological replicates (n = 3). WT and *lacI*$^-$ strains grown separately in mixture (2:1 v/v) of the specified carbon source and lactose minimal media. Cells were grown to exponential phase in the specified carbon source and diluted to an $OD_{600}$ = 0.01 at $t$ = 0. **(H)** Colony forming unit (CFU) counts of WT and *lacI*$^-$ strains. WT and *lacI*$^-$ strains grown in glycerol minimal media to exponential phase and plated on either LB or M9 minimal media supplemented with 0.4% lactose plates. Individual biological replicates (n = 3) ± SEM are shown in circles. **(I)** CFU and $OD_{600}$ measurements of *lacI*$^-$ strain during catabolite hyperflux lag and recovery phase. Cells were grown to exponential phase in M9 minimal media supplemented with 0.2% glycerol and diluted into M9 minimal media with 0.4% lactose to an $OD_{600}$ = 0.01 at $t$ = 0. Samples were plated on either LB or M9 minimal media supplemented with 0.4% lactose plates. Mean of biological replicates (n = 3) ± SEM of CFUs (circles) and $OD_{600}$ (squares) are shown.

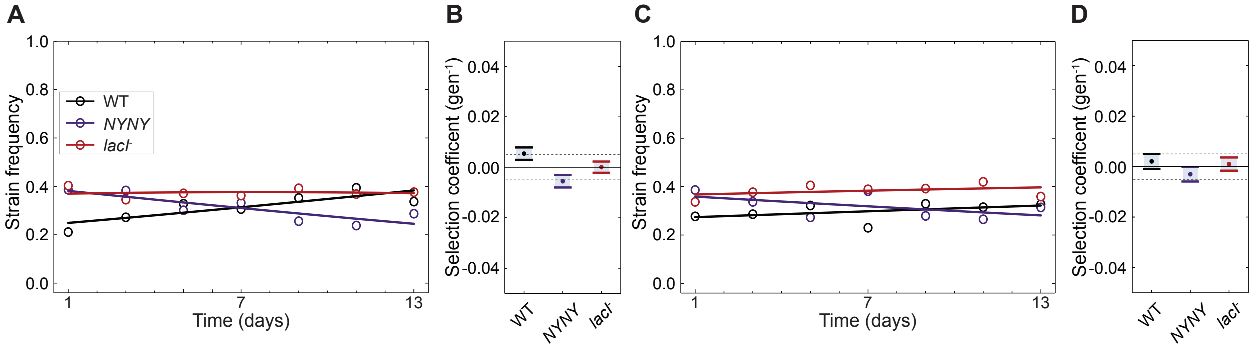


**Fig. S7. Controls for competition experiments selecting for or against gene regulation.**
WT, *lacI*⁻, and *NYNY* strains were combined in equal ratios based on normalized $OD_{600}$ and grown in either **(A)** glucose-only or **(C)** glycerol-only minimal media. Strains were diluted 1:100 into fresh media every 24 h for two weeks. Samples for analysis were taken in the same time intervals as in Fig. 3A,C. Relative exponential growth model was fit (solid lines) to infer selection coefficients. **(B,D)** Relative selection coefficients (points) and 95% confidence intervals (bars) are shown for each strain inferred from model fitting in (A,C) respectively. Dashed lines indicate upper and lower bound of confidence intervals inferred from competition in constant glycerol.

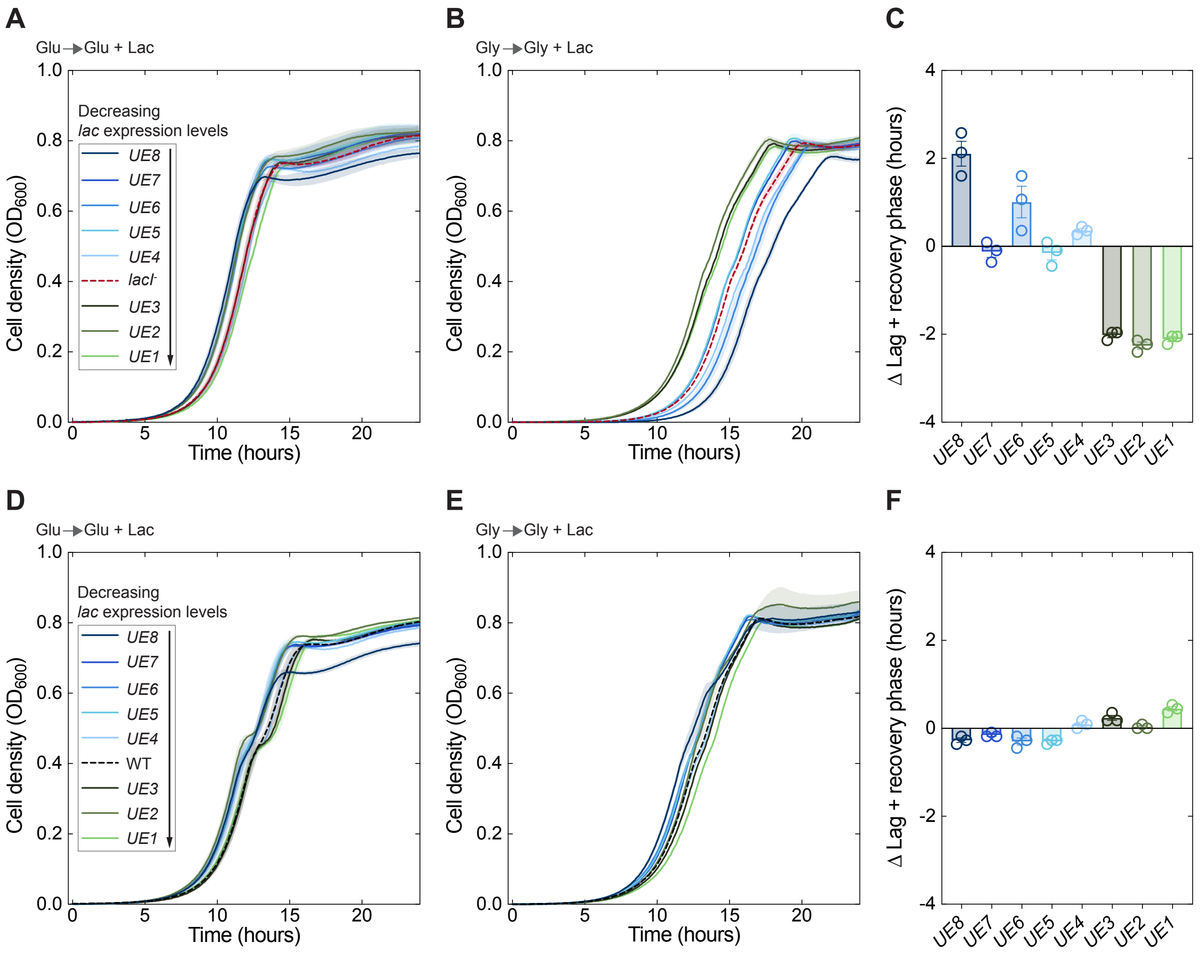


**Fig. S8. Additional data and analysis for epistatic interaction between sensing and control of gene expression.**

Diauxic growth dynamics of the UE strain library in *lacI*$^-$ **(A**,**B)** and WT **(D**,**E)** backgrounds. Each strain was grown separately in a mixture (2:1 v/v) of glucose (A,D) or glycerol (B,E) and lactose minimal media. Cells were grown overnight in glucose (A,D) or glycerol (B,E) and diluted to an $OD_{600}$ = 0.01 at *t* = 0. Legend indicates strains ordered by *lac* operon expression levels. Curves show mean $OD_{600}$ measurements taken every 5 minutes over biological replicates (n = 3); ± SEM is shown in shaded envelope. **(C**,**F)** Difference (Δ) in duration of lag and recovery phase during diauxic growth on glycerol and lactose (B,E) between each *lacI*$^-_{UE}$ strain and *lacI*$^-$ (C) or between each $WT_{UE}$ strain and WT (F). Time to $OD_{600}$ = 0.02 is shown measuring catabolite hyperflux lag and recovery (C) and gene induction lag and recovery (F). Variance of lag and recovery phase durations was calculated for each plot, yielding Var(Δ) = 2.31 $h^2$ (C) or 0.07 $h^2$ (F), which were found to be significantly different (F-test, *p*-value = 1.5e-12). Spearman's correlation coefficient (ρ) was calculated between Δ values and *lac* expression levels over UE strains, ρ = -0.86, *p*-value = 0.006 (C) and ρ = 0.73, *p*-value = 0.03 (F).

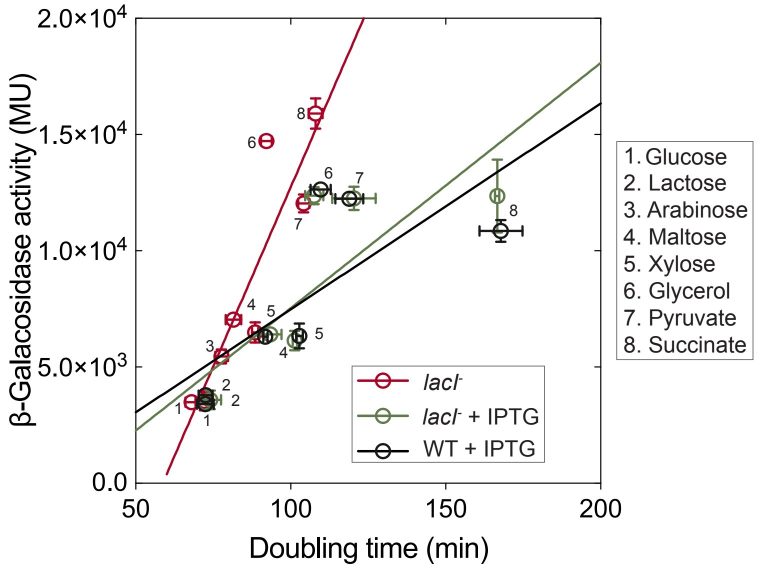


**Fig. S9. Relationship between β-galactosidase expression level and doubling time.**
Doubling time as a function of β-galactosidase activity levels in Miller units (MU) of the WT and *lacI*$^-$ from Fig. S5A,B. Numbers correspond to the single carbon sources used in the legend. Arabinose was omitted from the analysis for the WT and *lacI*$^-$ when supplemented with IPTG (see supplementary text for discussion). A linear relationship between β-galactosidase and growth rate was previously determined [56, 57], however the cost that is associated with the movement of IPTG through LacY was not accounted for [57]. We found that the linear relationship observed for *lacI*$^-$ without IPTG deviates from that of WT and *lacI*$^-$ strains with IPTG.

#### Supplementary Tables

**Table S1. Growth rates and β-galactosidase activity levels of *lacZ* ssrA-tagged *Escherichia coli* strains**

| | M9 Lactose | | M9 Lactose + 1% Casamino acids | | M9 Glucose | |
|---|---|---|---|---|---|---|
| Strain[a] | Growth rate, $h^{-1}$ ($\mu \pm \sigma$)[b] | β-Galactosidase activity, (MU ± $\sigma$) | Growth rate, $h^{-1}$ ($\mu \pm \sigma$)[b] | β-Galactosidase activity, (MU ± $\sigma$) | Growth rate, $h^{-1}$ ($\mu \pm \sigma$)[b] | β-Galactosidase activity, (MU ± $\sigma$) |
| WT | 0.55 ± 0.02 | 3050 ± 60 | 1.01 ± 0.05 | 3250 ± 120 | 0.64 ± 0.01 | N/A |
| *No Deg* | 0.54 ± 0.02 | 3040 ± 50 | 1.06 ± 0.01 | 3100 ± 50 | 0.62 ± 0.01 | N/A |
| *LAD* | 0.55 ± 0.02 | 2770 ± 100 | 1.03 ± 0.04 | 2770 ± 350 | 0.63 ± 0.01 | N/A |
| *AAV* | 0.56 ± 0.02 | 2730 ± 50 | 1.06 ± 0.03 | 2490 ± 110 | 0.63 ± 0.02 | N/A |
| *ssrA* | 0.56 ± 0.02 | 2430 ± 240 | 1.06 ± 0.03 | 2380 ± 40 | 0.63 ± 0.02 | N/A |
| *GSNY* | 0.55 ± 0.03 | 2140 ± 20 | 1.04 ± 0.04 | 2220 ± 20 | 0.63 ± 0.01 | N/A |
| *NYNY* | 0.54 ± 0.02 | 2400 ± 110 | 1.05 ± 0.03 | 2150 ± 200 | 0.64 ± 0.01 | N/A |

[a] Strains harboring addition of unique ssrA degradation tags to the 3' end of the *lacZ* gene in the chromosome. For more information regarding the *lacZssrA* strain library see Table S5 and [33].

[b] One-way ANOVA analysis showed no statistical significance of the mean growth rate (n ≥ 3) within each condition among the strains ($p$-value > 0.05).

**Table S2. Growth rates and β-galactosidase activity levels of the WT and *lacI⁻* strain in different carbon sources supplemented with IPTG and without**

| WT | | | | |
|---|---|---|---|---|
| | (-) IPTG | | (+) IPTG | |
| Carbon source | Growth rate, $h^{-1}$ ($\mu \pm \sigma$)[a] | β-Galactosidase activity, (MU ± $\sigma$)[b] | Growth rate, $h^{-1}$ ($\mu \pm \sigma$)[a] | β-Galactosidase activity, (MU ± $\sigma$)[b] |
| Glucose | 0.61 ± 0.01 | N/A | 0.57 ± 0.02 | 3410 ± 140 |
| Lactose | 0.57 ± 0.01 | 3120 ± 170 | 0.57 ± 0.01 | 3800 ± 230 |
| Arabinose | 0.53 ± 0.02 | N/A | 0.22 ± 0.01 | 7430 ± 270 |
| Maltose | 0.51 ± 0.01 | N/A | 0.45 ± 0.005 | 6310 ± 280 |
| Xylose | 0.47 ± 0.005 | N/A | 0.40 ± 0.005 | 6350 ± 530 |
| Glycerol | 0.44 ± 0.005 | N/A | 0.38 ± 0.01 | 12600 ± 160 |
| Pyruvate | 0.40 ± 0.005 | N/A | 0.35 ± 0.01 | 12200 ± 270 |
| Succinate | 0.38 ± 0.01 | N/A | 0.25 ± 0.01 | 10900 ± 460 |

| *lacI⁻* | | | | |
|---|---|---|---|---|
| | (-) IPTG | | (+) IPTG | |
| Carbon source | Growth rate, $h^{-1}$ ($\mu \pm \sigma$)[a] | β-Galactosidase activity, (MU ± $\sigma$)[b] | Growth rate, $h^{-1}$ ($\mu \pm \sigma$)[a] | β-Galactosidase activity, (MU ± $\sigma$)[b] |
| Glucose | 0.61 ± 0.02 | 3480 ± 270 | 0.57 ± 0.02 | 3460 ± 260 |
| Lactose | 0.58 ± 0.02 | 3550 ± 410 | 0.56 ± 0.03 | 3590 ± 410 |
| Arabinose | 0.54 ± 0.01 | 5460 ± 300 | 0.22 ± 0.01 | 6170 ± 640 |
| Maltose | 0.51 ± 0.02 | 7050 ± 260 | 0.45 ± 0.02 | 6400 ± 270 |
| Xylose | 0.47 ± 0.005 | 6480 ± 440 | 0.41 ± 0.005 | 6140 ± 420 |
| Glycerol | 0.45 ± 0.01 | 14700 ± 180 | 0.39 ± 0.01 | 12400 ± 370 |
| Pyruvate | 0.40 ± 0.01 | 12000 ± 400 | 0.35 ± 0.02 | 12300 ± 500 |
| Succinate | 0.39 ± 0.01 | 15900 ± 650 | 0.25 ± 0.001 | 12400 ± 1600 |

[a] Growth rates were calculated for the WT and *lacI⁻* strain in each carbon source (n ≥ 3).

[b] β-Galactosidase activity was measured in Miller units (MU) for the WT and *lacI⁻* in each carbon source (n ≥ 3). In the absence of IPTG, values were below or at background level except when WT cells were grown in lactose.

**Table S3. UP-element strains' modified sequence, β-galactosidase activity levels, and growth rates.**

| Strain name | Sequence | % change in β-gal activity[a] | Growth rate[b], $h^{-1}$ ($\mu \pm \sigma$) | | |
|---|---|---|---|---|---|
| | | | Glucose | Lactose | Glycerol |
| $WT_{UE1}$ | ATTGGACGT | 54% ± 3.9% | 0.62 ± 0.005 | 0.58 ± 0.03 | 0.44 ± 0.01 |
| $WT_{UE2}$ | GGTTTGAGC | 61% ± 4.6% | 0.63 ± 0.01 | 0.62 ± 0.06 | 0.46 ± 0.005 |
| $WT_{UE3}$ | ACCAGTAAG | 65% ± 3.6% | 0.63 ± 0.01 | 0.58 ± 0.04 | 0.45 ± 0.005 |
| WT | AGGCACCCC | 100% ± 7.8% | 0.62 ± 0.01 | 0.60 ± 0.02 | 0.45 ± 0.01 |
| $WT_{UE4}$ | GGATTAACG | 105% ± 7.8% | 0.60 ± 0.01 | 0.60 ± 0.03 | 0.44 ± 0.01 |
| $WT_{UE5}$ | ATAAACGCC | 112% ± 6.4% | 0.63 ± 0.005 | 0.64 ± 0.06 | 0.46 ± 0.01 |
| $WT_{UE6}$ | ACGCCTATC | 113% ± 9.2% | 0.63 ± 0.01 | 0.63 ± 0.05 | 0.45 ± 0.01 |
| $WT_{UE7}$ | TTCTTTCAG | 113% ± 12% | 0.63 ± 0.005 | 0.64 ± 0.04 | 0.45 ± 0.005 |
| $WT_{UE8}$ | CGGAAAGCG | 154% ± 14% | 0.64 ± 0.01 | 0.63 ± 0.04 | 0.47 ± 0.01 |
| $lacI^-_{UE1}$ | ATTGGACGT | 51% ± 7.8% | 0.62 ± 0.01 | 0.57 ± 0.04 | 0.45 ± 0.005 |
| $lacI^-_{UE2}$ | GGTTTGAGC | 60% ± 13% | 0.63 ± 0.02 | 0.64 ± 0.04 | 0.45 ± 0.005 |
| $lacI^-_{UE3}$ | ACCAGTAAG | 63% ± 4.7% | 0.62 ± 0.02 | 0.59 ± 0.04 | 0.44 ± 0.005 |
| $lacI^-$ | AGGCACCCC | 100% ± 9.4% | 0.61 ± 0.01 | 0.59 ± 0.01 | 0.45 ± 0.02 |
| $lacI^-_{UE4}$ | GGATTAACG | 104% ± 11% | 0.60 ± 0.01 | 0.60 ± 0.03 | 0.45 ± 0.005 |
| $lacI^-_{UE5}$ | ATAAACGCC | 110% ± 16% | 0.64 ± 0.02 | 0.69 ± 0.02 | 0.45 ± 0.005 |
| $lacI^-_{UE6}$ | ACGCCTATC | 115% ± 8.2% | 0.63 ± 0.01 | 0.64 ± 0.02 | 0.45 ± 0.01 |
| $lacI^-_{UE7}$ | TTCTTTCAG | 115% ± 16% | 0.63 ± 0.01 | 0.66 ± 0.03 | 0.45 ± 0.005 |
| $lacI^-_{UE8}$ | CGGAAAGCG | 146% ± 13% | 0.63 ± 0.02 | 0.67 ± 0.04 | 0.46 ± 0.005 |

[a] Average β-galactosidase activity levels compared to each respective parental strain (WT or *lacI*⁻). β-Galactosidase activity was measured using the Miller assay for each of the UP-element strains in glucose, lactose, or glycerol, and expressed as a ratio relative to the measured level of the respective parental strain (WT or *lacI*⁻) in each condition. The reported value is the average ratio over the three growth conditions. In glucose and glycerol conditions, cultures were supplemented with IPTG. β-Galactosidase activity was calculated from the mean (n = 3) β-galactosidase activity in Miller units (MU). Strains are numbered (1-8) in increasing β-galactosidase activity levels.

[b] Growth rates were calculated for the WT and *lacI*⁻ UE strain libraries in each carbon source (n ≥ 3).

**Table S4. Mutational landscape across phenotypic memory selection experiments**

| Coordinate[a] | Locus | Type | Mutation | Condition | Clones |
|---|---|---|---|---|---|
| 4174041 | *birA* | SNP | A -> T | Lac, 1:100 | 6/6 |
| 3641598 | *dtpB* | SNP | C -> T | Lac, 1:100 | 1/6 |
| 3827020 | *recG* | Insertion | IS1F element | Lac, 1:100 | 3/6 |
| 4182958 | *rpoB* | SNP | A -> G | Glu, 1:10,000 | 3/3 |
| 569919 | *ybcK* | SNP | G -> A | Lac, 1:10,000 | 1/3 |
| 3827020 | *recG* | Insertion | IS1F element | Lac, 1:10,000 | 3/3 |
| 322831 | *ykgF* | SNP | T -> A | G/L, 1:10,000 | 3/3 |
| 3815825 | Intergenic | SNP | C -> T | G/L, 1:10,000 | 3/3 |

[a] Coordinates are given for whole-genome sequencing data aligned to the MG1655 GCF_000005845.2 (RefSeq NC_000913.3) genome from NCBI.

#### Supplementary Movies 1,2 Captions

**Movies 1,2. WT and *lacI*$^{-}$ strains in periodic glucose-lactose fluctuations using microfluidics.** Growth of WT (Movie 1) or *lacI*$^{-}$ (Movie 2) in chemoflux microfluidics device. Cells were grown in minimal media with 0.4% lactose or 0.2% glucose as carbon source, and supplemented with 1% casamino acids. See also Fig. 1A.
